# Inter-individual variability in immune responses to AAV-mediated ocular gene delivery across species impedes reliable immunomonitoring profile

**DOI:** 10.1101/2025.06.02.656863

**Authors:** Duohao Ren, Gaelle Chauveau, Julie Vendomele, Emilie Cabon, Audrey Pineiro, Catherine Vignal-Clermont, Hanadi Saliba, Giuseppe Ronzitti, Anne Galy, Deniz Dalkara, Juliette Pulman, Divya Ail, Sylvain Fisson

## Abstract

Adeno-associated viruses (AAVs) have been used in gene therapy, especially for inherited retinal diseases. Despite their effectiveness in gene transduction, immune responses to the AAV capsid and transgene products have been reported, which can compromise both the efficacy and safety of AAV-mediated therapies. The eye is regarded as an immune-privileged organ where immune activity is constitutively suppressed. Here, we highlight that immunomonitoring in an ocular gene transfer reveals variable immune responses, whatever the species (human clinical trial, non-human primates, mice), the site of injection, the cassette, and the dose. We further explored factors contributing to this variability, investigating the correlation among immune parameters in a controlled experimental setting. In a syngeneic murine model after an intraocular injection of AAV, our results highlight an inter-individual variability of immune parameters, emphasizing the importance of considering inherent variability among individuals while designing personalized therapies.

## INTRODUCTION

Adeno-associated virus (AAV) vectors have become one of the most promising tools in gene therapy, largely due to their ability to deliver genetic material efficiently and safely into a variety of cell types. Unlike other viral vectors, AAV is non-pathogenic and exhibits minimal toxicity, making it suitable for treating genetic disorders such as hemophilia, muscular dystrophy, and retinal diseases(Sahel & Dalkara, 2019; Jauze *et al*, 2024; Yamaguti-Hayakawa & Ozelo, 2022; Saad *et al*, 2024). Since the first attempt of gene transfer in the retina with AAV in 1996 proving the efficiency of transduction(Ali *et al*, 1996), preclinical research in the application of AAV-mediated gene therapy for retinal diseases is on the rise. In 2007, the first clinical trial using AAV to treat Leber congenital amaurosis was initiated(Hauswirth *et al*, 2008) which eventually led to the first FDA-approved AAV-based ocular gene therapy, Luxturna (voretigene neparvovec), approved in 2017 for RPE65 mutation-associated retinal dystrophy(LUXTURNA® (voretigene neparvovec-rzyl) - Inherited Retinal Disease). And by the end of 2023, 94 clinical trials have been conducted or are ongoing targeting 12 retinal diseases with 6 AAV serotypes (clinicaltrials.gov, Key word: AAV, retina). These outputs have provided convincing proof for the application of AAV-based gene therapies to treat retinal genetic diseases. However, an often overlooked or under-reported aspect of AAV-based therapies, especially in the ocular field, is the immune responses induced by the AAV capsid and transgene product. The therapeutic success of AAV-based gene therapy is significantly influenced by the host’s immune response, which can on one hand limit the effectiveness of the treatment(Emami *et al*, 2023), and on the other cause adverse secondary effects such as inflammation.

Although the eye is considered as an immune-privileged organ(Streilein, 1999), safety of ocular gene therapy mediated by AAV is not guaranteed. Indeed, studies have shown that microglial cells can be activated after subretinal injection of AAVs in murine models(Khabou *et al*, 2018). Systemic humoral and cellular immune responses can also be elicited by AAV and transgene products that are delivered by subretinal injections(Ail *et al*, 2022; Vendomèle *et al*, 2024). A study evaluating the immune responses to AAV delivery in the retina of non-human primates (NHP) reported elevated levels of anti-AAV antibodies in the serum (systemic immune response) as well as ocular inflammation (local immune response)(Ail *et al*, 2022; Hakim *et al*, 2021). Another study, evaluating the cellular immune response by immunostaining in NHPs after AAV subretinal injections in the retina, reported CD8^+^ T-cell infiltration(Reichel *et al*, 2017).

Immune responses have not only been observed and reported in animal models used in gene therapy research, but also in clinical trials. Ocular inflammation was reported in patients undergoing ocular AAV gene therapy, and both anti-capsid humoral and cellular immune responses were reported, indicating the potential strong side effects of the AAV gene therapy(Bainbridge *et al*, 2008, 2015). Often, patients in these trials are provided with immunosuppression regimens that aim at managing the immune response-related side effects(Newman *et al*, 2021; Russell *et al*, 2017) Despite this, some patients developed ocular inflammation and systemic immune responses(Duke *et al*, 2022; Cehajic-Kapetanovic *et al*, 2020). An intriguing point is that non-consistent immune responses were observed whatever the condition in clinical trial. For example, in clinical trial NCT00749957, 3 out of 12 patients developed ocular inflammation and 5 out of 12 patients developed anti-capsid antibodies while no patients developed anti-capsid T cell response(Weleber *et al*, 2016). This has been supposedly attributed to differences in disease stage, treatment prior to gene therapy, genetics (Tong & Fairfax, 2020; Klein & Flanagan, 2016), lifestyle and environment(Huang *et al*, 2023) which are confounding factors that can influence the correlations among the immunomonitoring parameters in the patients and the efficiency of the gene therapy. Thus, we would like to test that in controlled animal models. Moreover, it can be considered as an added value to demonstrate that the same kind of immune response variability can be found whatever the species (human clinical trial, non-human primates, mice), the site of injection, the cassette, and the dose.

In the present study, we first highlighted the inter-individual differences in the anti-AAV immune responses in an ocular gene therapy clinical trial and in an NHP experiment. Next, we conducted analyses of local and systemic immune responses induced by AAV injection, maintaining consistent biological and experimental parameters to identify potential correlations among immune parameters. To that end, we administered AAV vectors through subretinal delivery in syngeneic mice and evaluated various immune parameters, including antibody production, T cell response, local inflammation, and cytokine secretion. Despite the controlled conditions, we observed significant inter-individual variability in immune responses, with only limited correlations identified among the assessed parameters.

## RESULTS

### Inter-individual variability of immune response is observed in the human clinical trial and NHP model after intraocular AAV gene transfer

Immunomonitoring after intraocular AAV2 gene transfer showed immune responses in both a human clinical trial(Bouquet *et al*, 2019) and NHPs(Ail *et al*, 2022). Since different strategies yielded distinct outcomes, we investigated the potential relations between the different immunomonitoring data from a human clinical trial described previously (NCT02064569)(Bouquet *et al*, 2019) (Figure 1A). In this clinical trial, 15 patients diagnosed with ND4 Leber Hereditary Optic Neuropathy (LHON), were distributed into four cohorts and treated with an intravitreal injection of a recombinant AAV2 vector carrying the ND4 gene at 4 different doses. A composite global ocular inflammation score (OIS) was determined using 4 separate grades according to Standardization of Uveitis Nomenclature (SUN) classification (Figure 1B). In addition, total antibody (TAb) and neutralizing antibody (NAb) levels against the capsid were also measured by ELISA and NAb assays respectively in patients. TAb and NAb levels were generally increased in patients’ post-injection serum samples, with no clear dose effect (Figures 1C, 1C’). Immune profiles were generated for each patient, incorporating fold-change of TAb and Nab against AAV2, maximal OIS and anti-capsid cellular immune responses measured by IFNγ ELISpot in peripheral blood mononuclear cells (PBMCs) isolated from patients. The immunomonitoring revealed distinct patterns in patients who received the same dose, exhibiting different levels of ocular inflammation scores (OIS), cellular and humoral responses regardless of the AAV dose received (Figures 1D-G, Supplementary table 1).

**Figure 1.**
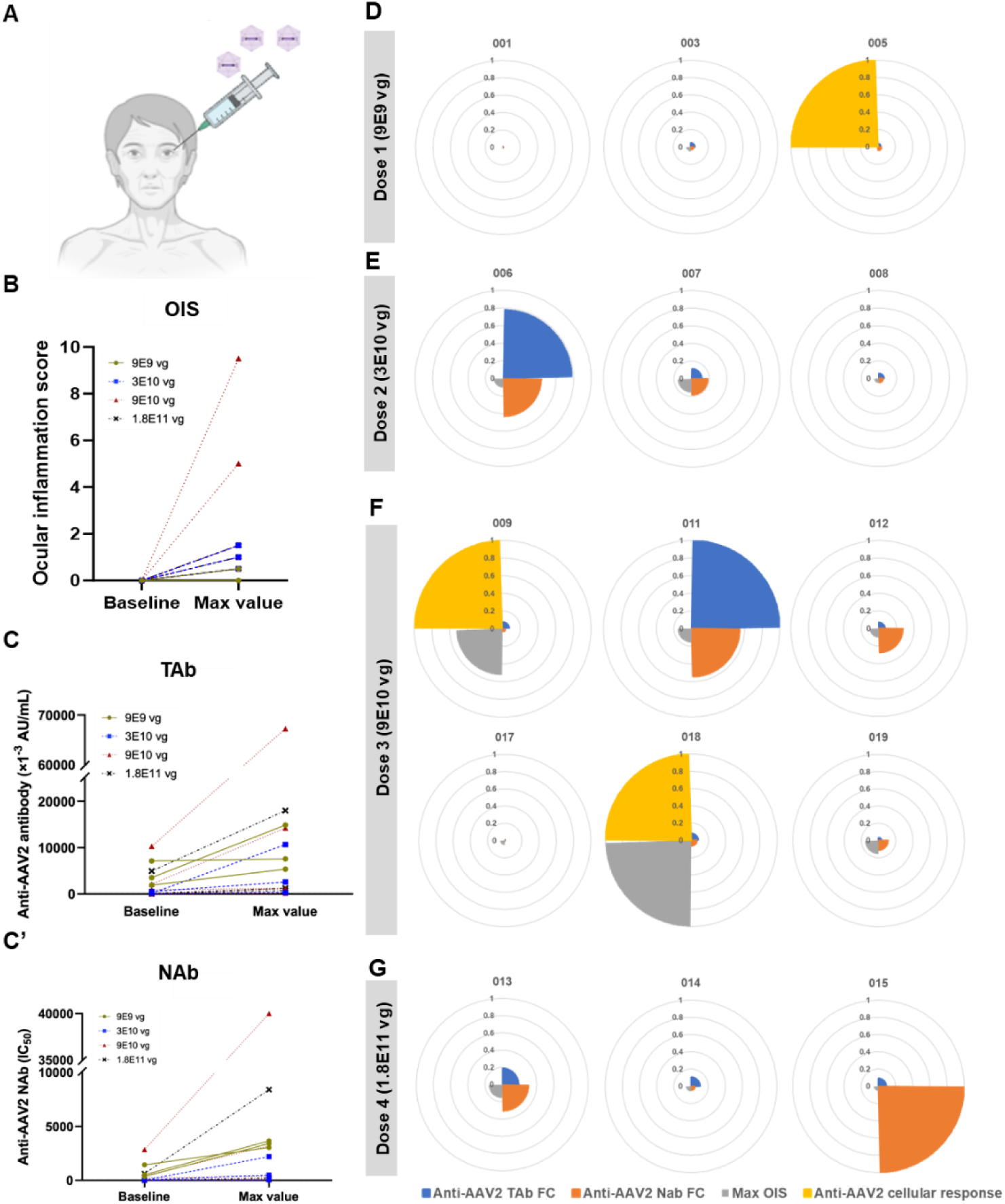
Inter-individual variability of immune response is observed in the human clinical trial (NCT02064569) after ocular AAV gene transfer. **A** Schematic of ocular gene injections in patients. **B** Ocular inflammation score (OIS) in human patients who received intraocular AAV injections at baseline and maximal value (obtained within 20 weeks after injection, except patient 003 for 40 weeks). **C, C’** Total antibody (TAb) (**C**) and neutralizing antibody (NAb) (**C’**) levels measured by ELISA and NAb assay against AAV2 in human patients who received intraocular AAV injections at baseline and maximal value (obtained within 20 weeks after injection). IC_50_: half maximal inhibitory concentration. **D-G** Immune profiles of individual patients, who received dose 1 – 9×10^9^ vg (**D**), dose 2– 3×10^10^ vg (**E**), dose 3– 9×10^10^ vg (**F**), dose 4– 1.8×10^11^ vg (**G**) of AAV, showing maximum ocular OIS,foldchange (FC) of anti-AAV2 TAb and NAb, anti-AAV2 cellular immune response. Each slice of the pie corresponds to one immune parameter which is normalized against the highest value for the parameter. The code of each patient is shown on the above each chart. vg: vector genome.

In order to further investigate immune responses under more controlled environmental conditions, immunomonitoring data from a study on NHPs were analyzed(Ail *et al*, 2022). All the 8 NHPs received an intravitreal injection of 5×10^11^ vector genomes (vg) of AAV2.7m8 encoding ChrimsonR in both eyes (Figure 2A). Ocular inflammation was assessed by slit lamp 1-month post-injection and scores were determined by SUN classification on anterior chamber cells (ACC), anterior chamber flare (ACF), vitreous haze (VH), and vitreous cells (VC). ELISA and NAb assays were used to measure capsid-specific TAb and NAb levels 2 to 3 months post-injection (PI). Both TAb and NAb against capsid in NHP sera increased after AAV administration (Figures 2B, 2B’). Normalized immune profiles of NHPs highlighted varying contributions of humoral immunity and inflammation by slit lamp, including ACC, ACF, VH, and VC to the overall response (Figure 2C, Supplementary table 2). Both species exhibited variability in immunomonitoring outputs following intraocular AAV injection.

**Figure 2.**
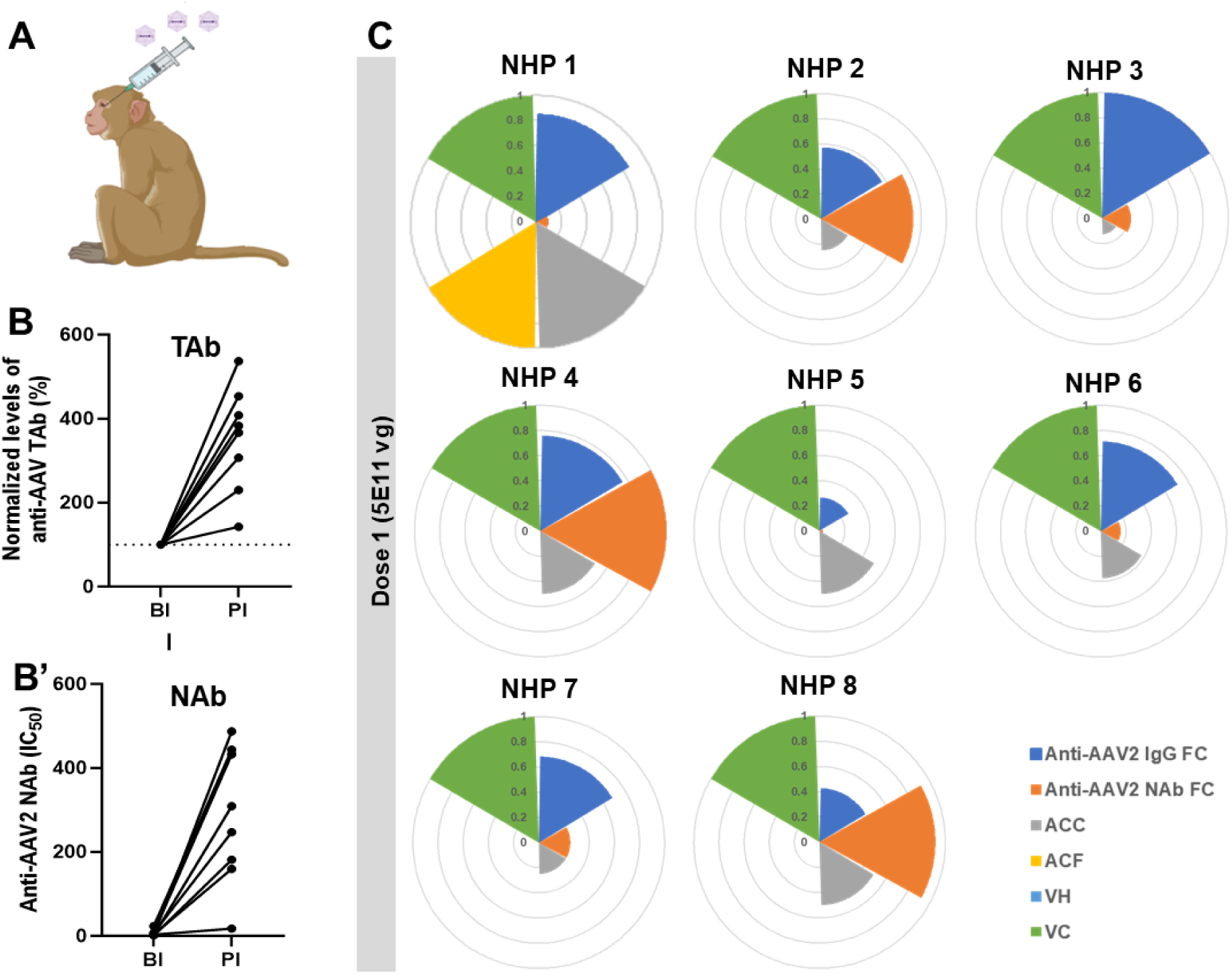
Inter-individual variability of immune response is observed in the NHP model after ocular AAV gene transfer. **A** Schematic of ocular gene injections in NHPs. **B, B’** Total antibody (TAb) **(B)** and neutralizing antibody (NAb) **(B’)** levels measured by ELISA and NAb assay against AAV2 in NHPs who received intraocular AAV injections with dose 1 – 5×10^11^ vg before (BI) and 2 to 3-month post injection (PI). IC_50_: half maximal inhibitory concentration. **C** Immune profiles of individual NHPs showing FC of anti-AAV2 TAb and NAb, grading score at month 1 for Anterior Chamber Cells (ACC), Anterior Chamber Flare (ACF), Vitreous Haze (VH) and Vitreous Cells (VC). Each slice of the pie corresponds to one immune parameter which is normalized against the highest value for the parameter. The code of each NHP is shown on the above each chart. Vg: vector genome.

### Systemic adaptive immune responses both against capsid and transgene product are induced after subretinal injections of AAV8-GFP-HY in mice

Due to the diversity in the disease stage, genome and environmental factors in humans and NHPs, syngeneic murine models seem to be pertinent to explore the consistency of the immune responses following AAV-mediated AAV gene transfer. Previous studies have demonstrated that AAV intraocular injections can induce both humoral and cellular immune responses against capsids and transgene products that can be detected systemically in blood or in lymphoid organs in murine models(Li *et al*, 2024; Ertl, 2021; Calcedo *et al*, 2013; Bucher *et al*, 2021); however, these studies typically focused on limited immune parameters like antibody production and local inflammation. To comprehensively assess the humoral and cellular immune responses against both capsids and transgene products, a male peptide named HY which was known to induce systemic T cell responses(Vendomèle *et al*, 2018, 2024) in female mice was packaged into AAV8 capsid along with GFP. The AAV vectors (5×10^10^ vg/eye) were administered into the eye of female C57Bl/6 mice via subretinal injection (Figure 3A). Sera and spleens were collected 21 days post-injection as previously described(Vendomèle *et al*, 2018, 2024) to analyze the cellular and humoral immune response against the capsid and transgene product (Figure 3A). IFNγ ELISpot revealed a significant increase in activated T cells against AAV (p-value = 0.0001) and transgene product (p-value < 0.0001) in AAV-injected mice compared to PBS-injected control mice (Figures 3B, 3C and Figure S1A). HY peptides (pHY) are composed of DBY, activating CD4^+^ T cells and UTY, activating CD8^+^ T cells(Carpentier *et al*, 2015). ELISpot assay showed an IFNγ secretion by CD4^+^ and CD8^+^ T cell specific to transgene product (Figures S1B, S1C). ELISA measurements showed that anti-AAV8 and anti-GFP antibodies significantly increased (p-value < 0.0001) after AAV subretinal injection compared to PBS-injected control mice (Figures 3D, 3E). The systemic cytokine profile contributing to the inflammation and immune response in mice was evaluated following AAV subretinal injection. Spleen cells were isolated from mice that received either PBS or AAV. These spleen cells were then stimulated *in vitro* with pHY or AAV. A cytometric bead array (CBA) was used to measure cytokines in the culture supernatant to determine the cell polarizations: Th1/Tc1 (IL-2, IFNγ, TNFα, GM-CSF), Th2/Tc2 (IL-4, IL-10, IL-13), Th17/Tc17 (IL-17), and those involved in inflammation and migration (IL-1β, IL-6, RANTES, MCP-1) in the culture supernatant to determine the cell polarizations. Radar charts were generated to visualize the proportional production of these cytokines. Spleen cells from PBS-injected control mice mainly did not secrete cytokines (Figures 4A, 4A’). In contrast, spleen cells from AAV-injected mice produced production of multiple cytokines (Figures 4B, 4B’). Among cytokines involving inflammation and migration, only RANTES (p-value = 0.01) showed a significant increase (Figure S2A). The Th1/Tc1 cytokines, IL-2 (p-value = 0.0046), IFNγ (p-value = 0.0001), TNFα (p-value = 0.0388), were significantly upregulated (Figure S2B), in response to *in vitro* AAV stimulation. Other cytokines remained unchanged (Figures S2A, S2B, S2C, and S2D). Upon *in vitro* stimulation with pHY, cytokines such as RANTES (p-value < 0.0001), IFNγ (p-value = 0.0001), TNFα (p-value = 0.0007), and IL-10 (p-value = 0.0024) were significantly upregulated (Figure S3).

**Figure 3.**
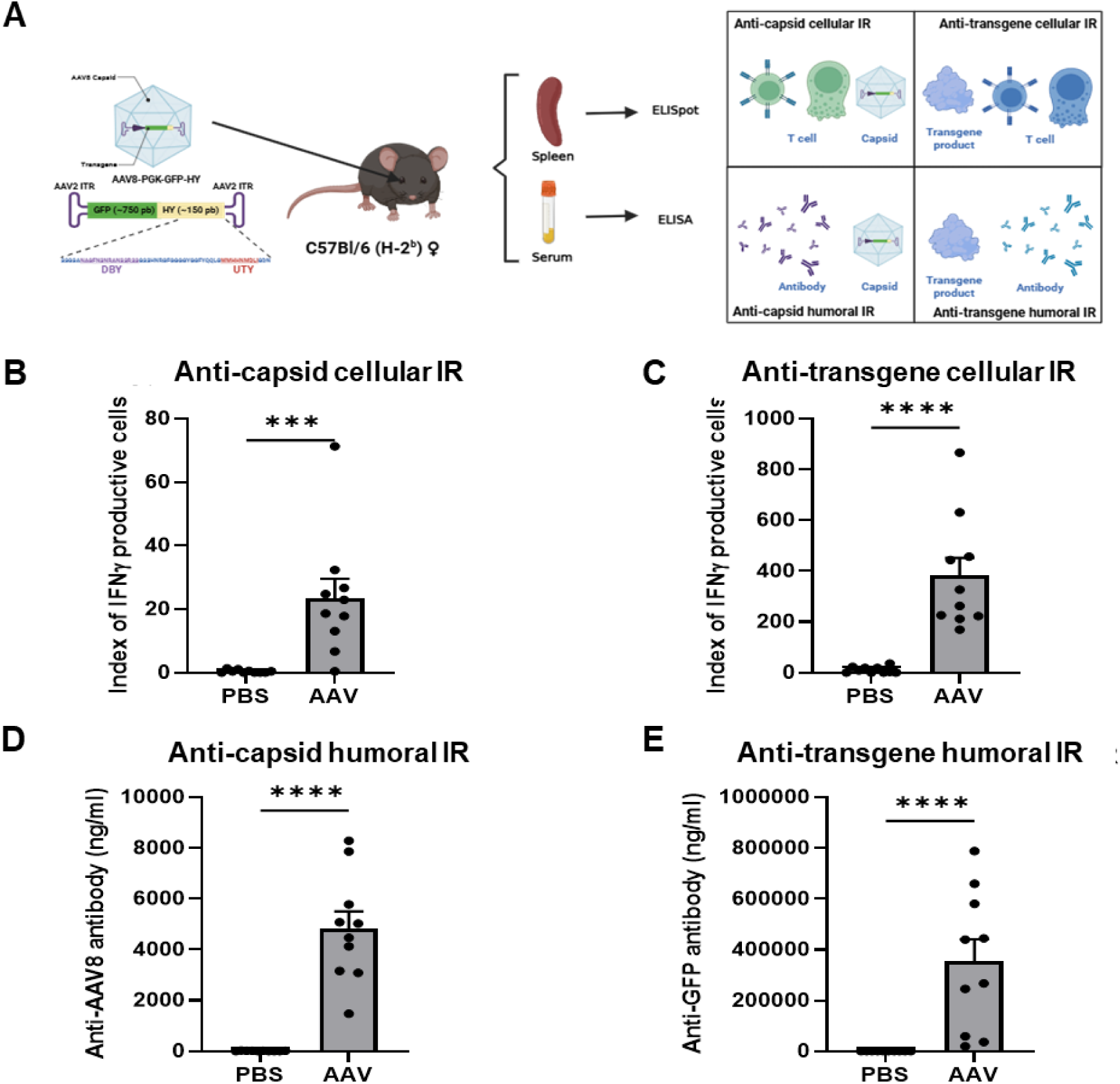
Systemic adaptive immune responses are induced after subretinal injections of AAV8-GFP-HY in mice. **A** Schematic representation of the experimental procedure showing the subretinal injection of AAV8-GFP-HY (5×10^10^ vg) in mouse, followed by harvest of the spleen and serum 21d post-injection (PI) to test cellular and humoral responses against the capsid and transgene. **B, C** T cell activation measured by ELISpot assay against **(B)** the AAV8 capsid and **(C)** the transgene-HY. **D, E** Antibody levels measured by ELISA against **(D)** the AAV8 capsid and **(E)** the transgene-GFP. Data information: Results obtained from 2 independent experiments (n=10 per group). Bars correspond to mean + SEM. *P < 0.05, **P < 0.01, ***P < 0.001, and ****P < 0.0001 with unpaired Mann-Whitney test.

**Figure 4.**
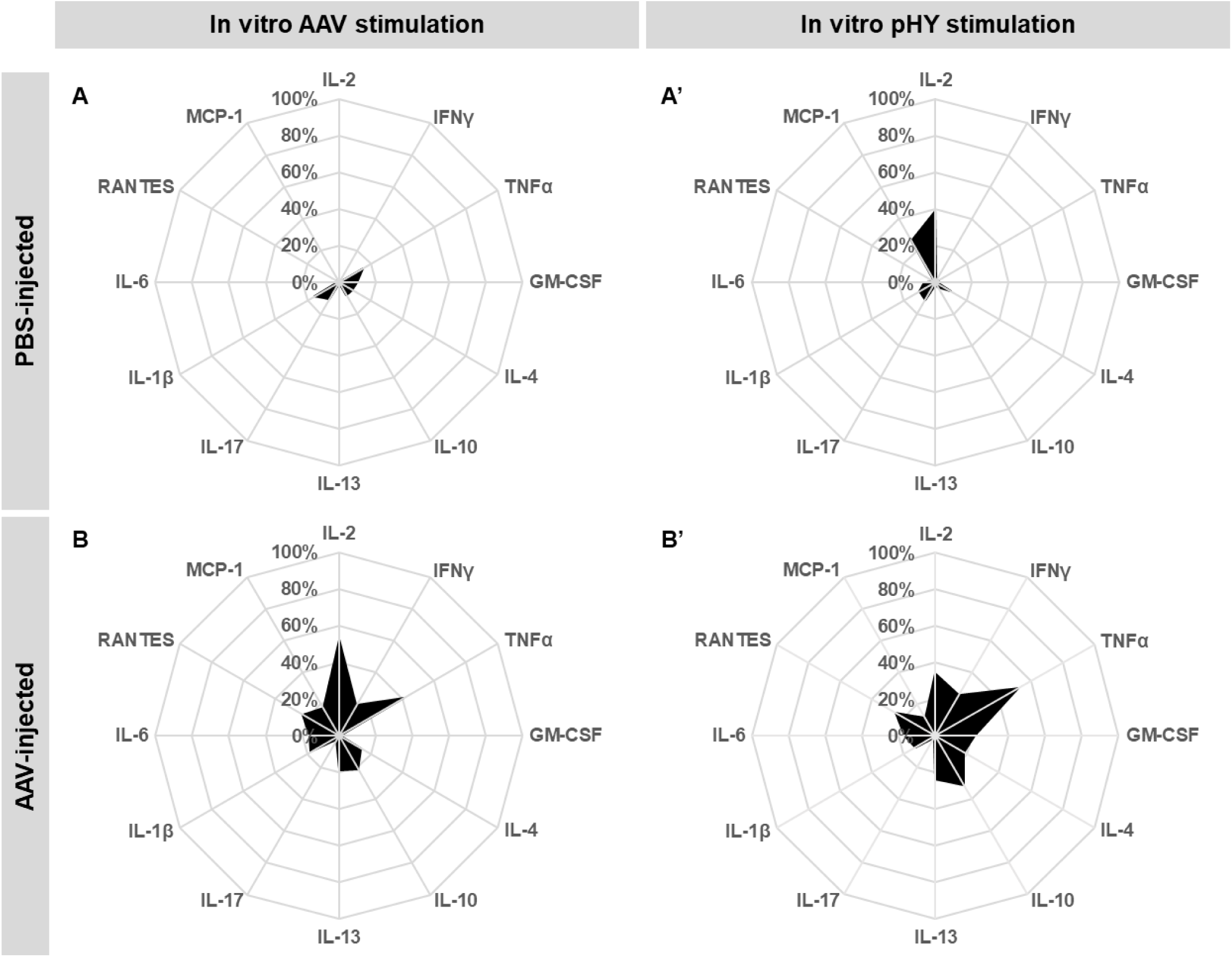
Global view of the systemic pro-inflammatory cytokine profile in mouse 21d post-injection (PI) of AAV-GFP-HY. **A, A’** Cytokine profiles of PBS-injected group with **(A)** AAV *in vitro* stimulation and **(A’)** peptide HY (pHY) stimulation. **B, B’** Cytokine profiles of AAV-injected group with **(B)** AAV *in vitro* stimulation and **(B’)** pHY stimulation. The 100% radar scale fits to the maximum value of cytokine secretion, and the black area values correspond to the mean of these cytokine secretions. Cytokines tested are ILs: Interleukin, TNFα: Tumor Necrosis Factor alpha, IFNγ: Interferon gamma, GM-CSF: Granulocyte-Macrophage Colony-Stimulating Factor, RANTES: Regulated upon Activation, Normal T cell Expressed and Secreted (also known as CCL5), MCP-1: Monocyte Chemoattractant Protein 1 (also known as CCL2). The concentration of each cytokine in each condition can be found in Supplementary figure 2, 3 and Supplementary table 3.

### Local transgene expression in the retina is correlated with cytotoxicity against transgene-expressing cells post-injection of AAV-GFP-HY

Further, the level of transgene expression was measured to assess the efficacy of AAV transduction and its relationship with cytotoxic effects using Droplet Digital PCR (ddPCR). In AAV-injected mice, significant expressions of the HY (p-value = 0.0014) and GFP (p-value = 0.0002) transgenes were observed 21 days after injection, and a correlation between the expression levels of both transgenes was identified (R² = 0.818) (Figures 5A, 5B). To evaluate the clearance ability of specific antigen-expressing cells in these mice, an *in vivo* cytotoxicity assay was performed using male spleen cells expressing the HY antigen as target cells which means the more target cells survive, the less cytotoxicity there is. We found a significant reduction (p-value = 0.0004) in the number of male target cells in AAV-injected mice, indicating the development of HY-specific cytotoxicity (Figure 5C). Direct correlations between target cell survival in the blood and retinal transgene expression (R² = 0.7702 for GFP; R² = 0.7935 for HY) were demonstrated suggesting an inverse correlation between cytotoxicity effect and transgene expression in the mice (Figures 5D, 5E).

**Figure 5.**
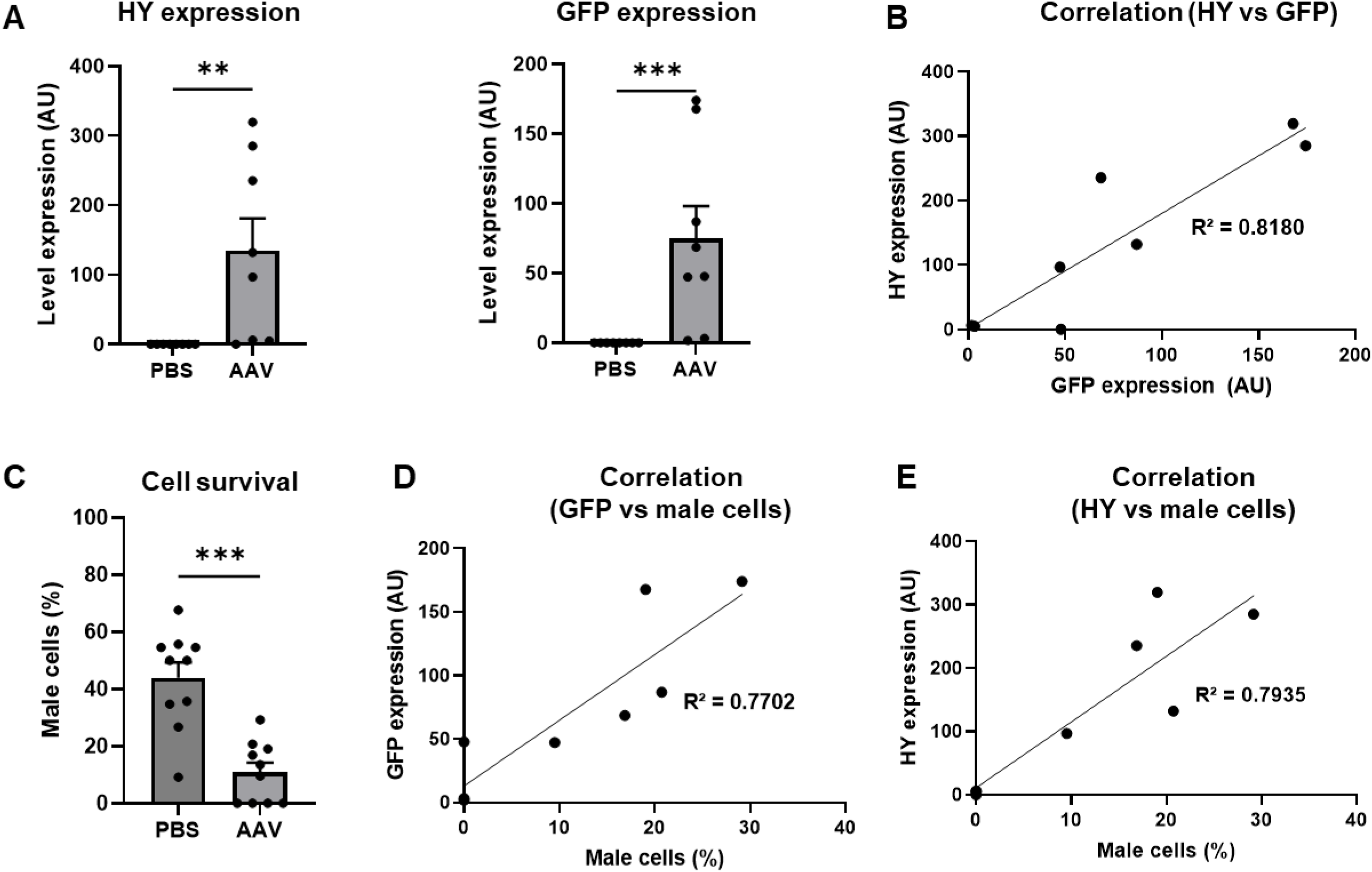
Local transgene expressions in the retina are correlated with cytotoxicity against transgene^+^ cells post-injection (PI) of AAV-GFP-HY. **A** HY and GFP transgene expression levels in female mouse retina 21d PI measured by ddPCR. AU: Arbitrary Units. **B** Correlation between HY and GFP expressions. **C** Activation of cytotoxic cells specific to the HY peptide 21 PI evaluated by *in vivo* cytotoxicity assay. **D, E** Correlation between the survival of male cells and local transgene expression in AAV-injected mice for **(D)** GFP and **(E)** HY. Data information: Results obtained from 2 independent experiments (n=8 per group). Bars correspond to mean + SEM. *P < 0.05, **P < 0.001, and ***P < 0.0001 with unpaired Mann-Whitney test. R^2^ is calculated via linear regression.

### Local inflammation and correlated expressions between MHC II molecules and CYBB are found in the retina post-injection of AAV-GFP-HY

The direct correlation between target cell survival and retinal transgene expression suggested a potential antigen presenting cell (APC) contribution to present transgene product peptides, associated with local inflammation. It was assessed by evaluating the transcript expression of MHC II molecules(Lipski *et al*, 2017) (H2-Eb1 and H2-Ab1) and the type 1 macrophage marker CYBB(Frazão *et al*, 2015) in the retina, using ddPCR 21 days after the injection. We observed a significant increase in the expression of both MHC II molecules (p-value = 0.0002) 21 days after the subretinal injection (Figure 6A). Since H2-Eb1 and H2-Ab1 are co-dominant molecules, their correlated expressions were confirmed (R² = 0.8686) (Figure 6B). Interestingly, the expression of CYBB increased significantly in AAV-injected mice (p-value = 0.0002) (Figure 6C), and correlations were found between CYBB and MHC II molecules expression (R² = 0.9659 for H2-Eb1; R² = 0.8165 for H2-Ab1), suggesting a link between antigen presentation and macrophage activation (Figures 6D, 6E).

**Figure 6.**
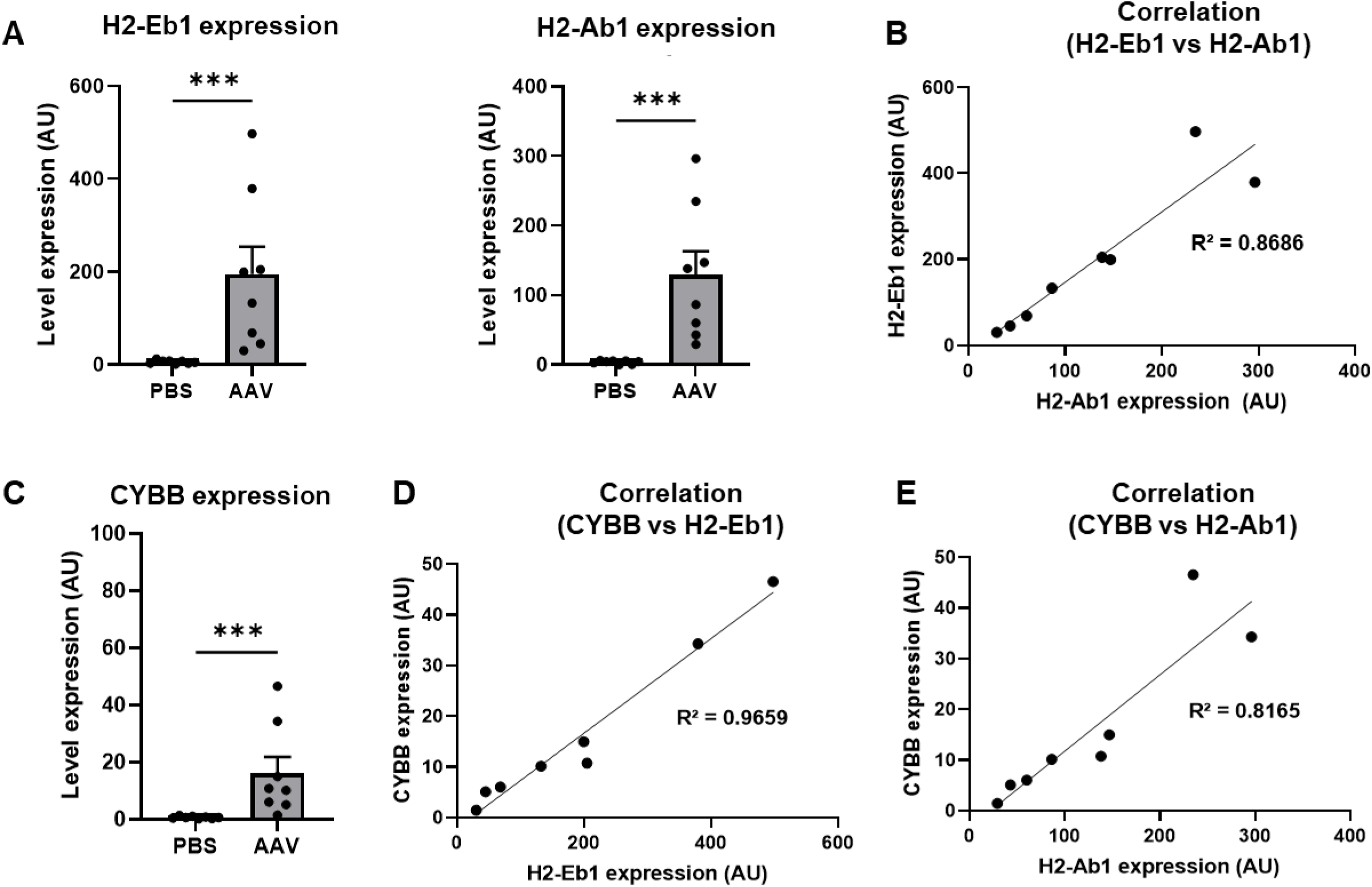
Local inflammation and correlated expressions between MHC II molecules and CYBB are found in the retina post-injection (PI) of AAV-GFP-HY. **A** Gene expression of H2-Eb1 and H2-Ab1 measured by ddPCR in the mouse retina 21d PI. AU: Arbitrary Units. **B** Correlation between H2-Eb1 and H2-Ab1 expression levels. **C** Gene expression of CYBB measured by ddPCR in the mouse retina 21d PI. **D, E** Correlation between **(D)** CYBB and H2-Eb1 expression; and **(E)** CYBB and H2-Ab1 expression. Data information: Results obtained from 2 independent experiments (n=8 per group). Bars correspond to mean + SEM. *P < 0.05, **P < 0.001, and ***P < 0.0001 with unpaired Mann-Whitney test. R^2^ is calculated via linear regression.

### Diversity in immune responses is noticed in syngeneic mice 21d post-injection of AAV-GFP-HY

After analysis of all immune parameters typically collected for immunomonitoring in clinical trials, variability in the immune parameters was observed even in this syngeneic model. To figure out the main factors driving this variability, principal component analysis (PCA) was performed, and angle sectors were then used to show the immune profiles and cytokine profiles of each mouse. In angle sectors, all analyzed transgene expression, immune parameters and significantly regulated cytokines were normalized to their corresponding highest values to provide a more direct visualization. The immune profile included transgene expression (HY-Tg and GFP-Tg), local inflammation (H2-Eb1, H2-Ab1 and CYBB), humoral immune response (AAV-Ab and GFP-Ab) and cellular immune response (%male, ELISpot-HY, ELISpot-DBY, ELISpot-UTY, ELISpot-AAV-PGK-Luc2 and ELISpot-AAV-PGK-GFP-HY). The cytokine profile was composed of cytokines significantly affected by AAV *in vitro* stimulation (IL-2, IFNγ, TNFα and RANTES) or HY *in vitro* stimulation (IFNγ, TNFα, IL-10 and RANTES). No similar immune profiles were found in AAV-injected mice suggesting that the immune responses against AAV and its transgene product may vary among mice (Figures 7A, 7B, Supplementary table 3). The Principal Component Analysis (PCA) results confirmed that AAV-injected mice could be clearly separated from PBS-injected mice as expected and that the AAV-injected group exhibited a broad distribution of the individuals which was consistent with immune and cytokine profiles, highlighting the variability of immune parameters after AAV subretinal injection (Figure 7C).

**Figure 7.**
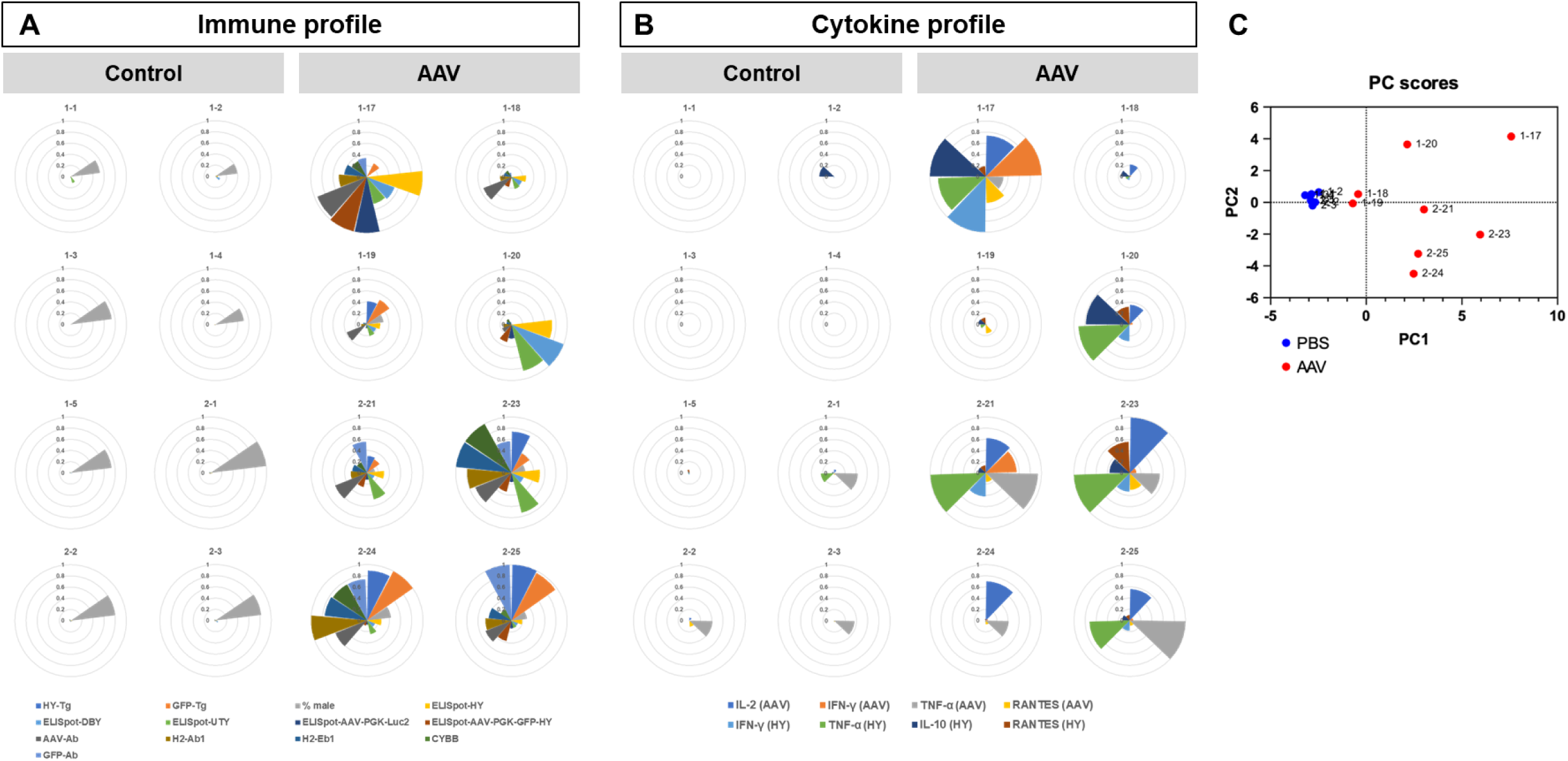
Diversity in immune responses is noticed in syngeneic mice 21d post-injection (PI) of AAV-GFP-HY. **A** Immune profiles of each individual mouse showing transgene expression (HY-Tg and GFP-Tg), local inflammation (H2-Eb1, H2-Ab1 and CYBB) and humoral immune response (AAV-Ab and GFP-Ab) and cellular immune response (%male, ELISpot-HY, ELISpot-DBY, ELISpot-UTY, ELISpot-AAV-PGK-Luc2 and ELISpot-AAV-PGK-GFP-HY) 21d PI of PBS (control) and AAV8-GFP-HY (AAV). **B** Cytokine profiles of individual mouse showing cytokine levels (AAV *in vitro* stimulation: IL-2, IFNγ, TNFα and RANTES; HY *in vitro* stimulation: IFNγ, TNFα, IL-10 and RANTES) 21d PI of PBS (control) and AAV8-GFP-HY (AAV). Each slice of the pie corresponds to one immune parameter which is normalized against the highest value into percentage for the parameter. **C** Principal component analysis of PBS-injected mice (blue dots) compared to AAV-injected mice (red dots). Data information: The mouse code is shown related to the dot. The code of mouse is shown on the above each chart, Results obtained from 2 independent experiments (n=8 per group).

### Absence of correlation between immune parameters in syngeneic mice 21 days following AAV-GFP-HY delivery

To explore relationships among immune parameters, a correlation analysis was conducted. With a correlation matrix, R² values for each parameter pair was displayed and was further visualized using a network analysis to illustrate the connections among these parameters. Distinct correlations emerged between certain parameters; for instance, anti-transgene product antibody levels showed a clear correlation with transgene expression, which in turn was linked inversely to male cell death, indicating the cytotoxic effect. However, most parameters displayed no significant correlations, particularly across the four primary domains: local transgene expression, local inflammation, systemic immune response, and cytokine profile (Figures 8A, 8B). The absence of correlation between immune markers suggests that subretinal AAV administration leads to individualized immune responses, with significant variation between subjects. This indicates that each subject may mount a distinct immunological reaction to the treatment.

**Figure 8.**
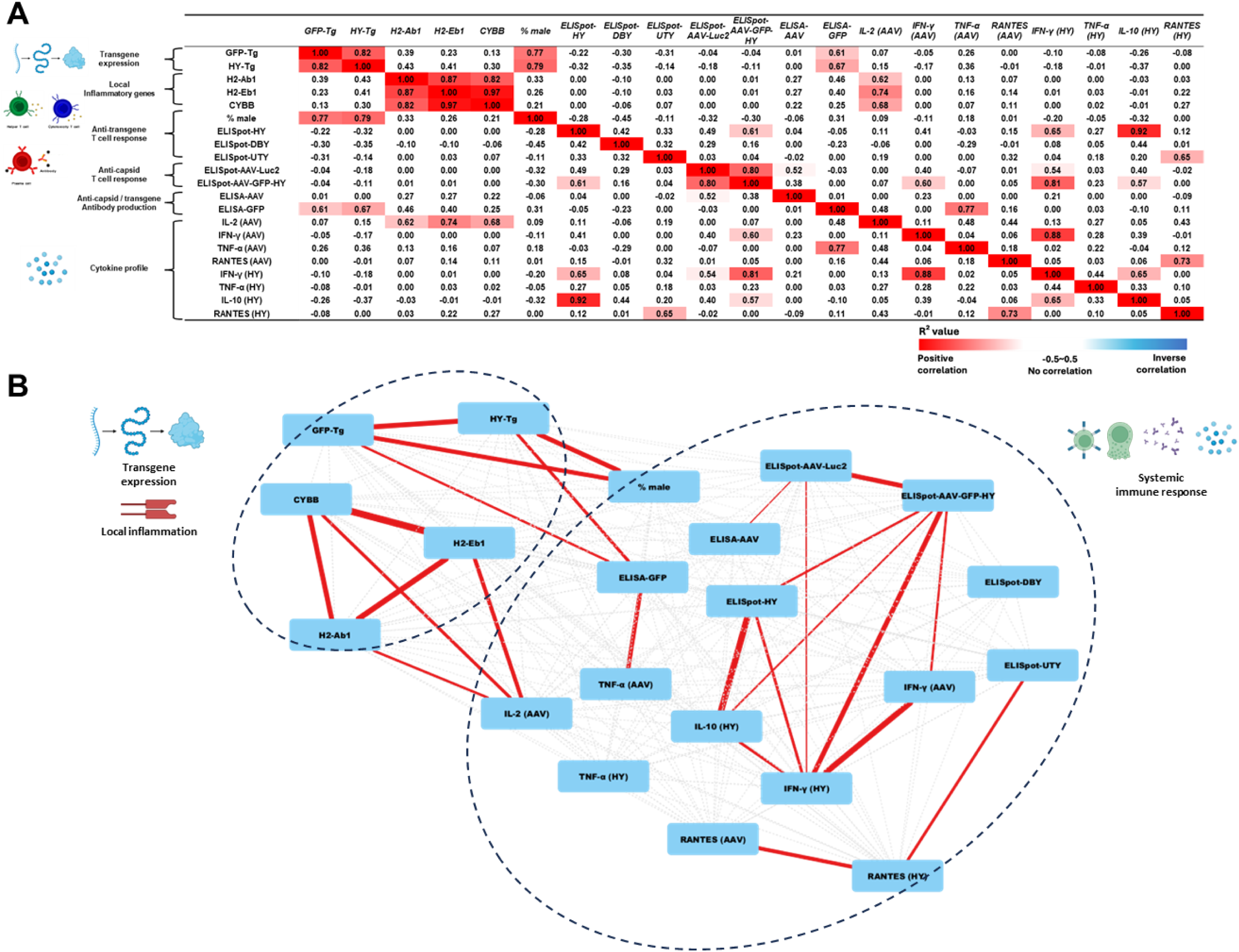
Immune parameters show no correlation in syngeneic mice 21 days following AAV-GFP-HY delivery. **A** Correlation matrix showing the coefficient of determination of each pair of immune parameters tested experimentally. Positive correlations are marked in shades of red with R² above 0 and negative correlations in shades of blue with minus label (−) before R². The intensity of shade corresponds to the strength of correlation. **B** Network analysis map generated based on the correlation matrix. The color of the lines connecting two parameters (edges) indicate positive (red), negative (blue) or no (grey) correlation and the thickness of the edges indicate the strength of correlation with thicker lines indicating higher correlation and dotted lines showing no correlation.

## DISCUSSION

Gene therapy offers a promising approach for treating genetic diseases, with AAV vectors employed to deliver the therapy (Niemeyer *et al*, 2009; Bainbridge *et al*, 2003; Miyoshi *et al*, 1997). Nevertheless, immune responses triggered by AAV capsids and transgene products remain a concern, as they are observed both peripherally and locally in preclinical and clinical studies, and they are generally considered as factors that can negatively affect the effectiveness of gene therapy(Ertl, 2022; Shen *et al*, 2022; Ren *et al*, 2022). We highlight here that immunomonitoring results in clinics vary among patients. The immune response variations observed in patients is usually attributed to the stage of disease, treatment prior to gene therapy, genetics, environment and lifestyle. However, in our study despite the use of a syngeneic murine model treated with AAV vectors in highly controlled conditions (same sex, same age, kept in the same environment, subjected to the same treatment), a considerable inter-individual variability was still observed, similar to the high inter-individual variability noticed both in human clinical and NHP data.

Inter-individual variability of immune response after intraocular AAV injections was evident in humans and NHPs. None of the individuals, whether human or NHP, who received the same dose and vector injection exhibited identical immune responses. This observation confirms that there is a widespread variability in heterogeneous individuals which has not been well understood and emphasizes the critical need to identify the factors contributing to this inter-individual variability. In order to confirm if the inter-individual variability persists despite the route of administration, transgene and serotypes, another AAV vector was applied in murine experiments. Thus, systemic immune response induced by AAV subretinal injection was characterized in the syngeneic murine model. HY-GFP were packaged into AAV which was previously shown to induce systemic T cell immune responses(Vendomèle *et al*, 2018, 2024) against transgene products. All the mice developed adaptive systemic immune responses and surprisingly innate immune responses persisted 3 weeks after the injection. In previous studies aiming to explore the factors impacting the AAV-induced immune responses, a dose-dependency was repetitively identified(Ronzitti *et al*, 2020; Hamilton & Wright, 2021) which can also be influenced by serotype (Gehrke *et al*, 2022; Bentler *et al*, 2023; Chan *et al*, 2021), administration route(Ge *et al*, 2001), transgene(Boisgerault *et al*, 2013; Xiong *et al*, 2019; Khabou *et al*, 2018; Pfromm *et al*, 2022) and the sex of the recipient(Clare *et al*, 2025). In the context of this study, optimization of vector dose and transgenes could help reduce the risk of inducing a potential immune response. Furthermore, developing novel AAV vectors with lower immunogenicity can be another alternative option to evade part of the immune responses directed against the capsid. In addition to directed evolution, capsid shuffling and rational design, modeling in silico and machine learning are increasingly being used for AAV capsid development. In silico strategies can predict which mutations are likely to enhance AAV functionality and narrow down potential candidates to test in experimental setting(Wec *et al*, 2021).

Correlation data analysis revealed a novel finding: very few significant relationships between immune parameters, including local transgene expression, local inflammation, and systemic immune responses. This relative independence of immune factors has not been previously documented. Among the studied parameters, transgene expression level appears to be inversely correlated with the cytotoxicity against transgene expressing cells. In addition, it also seems in our murine model that the humoral immune response against the transgene is correlated with the transgene product level of expression, but not with cytotoxicity. Further investigations, including other controls and transgene cassettes, should help to clarify if there is an impact of the transgene type on this absence of correlation between humoral and cellular adaptive immune responses. Interestingly, our results suggested a link between antigen presentation (MHC II) and macrophage activation (CYBB) in the retina. However, the absence of correlation found with the commonly used clinical immunomonitoring methods, such as ELISA and ELISpot(Bainbridge *et al*, 2015; VIGNAL-CLERMONT *et al*, 2023; Britten *et al*, 2013), which investigate peripheral adaptive immune responses, suggests that they may not fully capture the complexities of immune reactions in ocular gene therapy patients. Thus, the use of these immunomonitoring techniques may not reflect the actual immune status and cannot be used to determine the application of immunosuppression regimens in patients. It would also be informative in further studies in animal models to investigate the potential link between the immune response variability and the percentage of retinal area impacted by the injection, such as with RNA in-situ hybridization or immunofluorescence assays on retinal flatmounts, or a ddPCR assay to quantitatively detect AAV genome or transgene. Analysis of systemic cytokine profiles and putative inflammatory biomarkers(Calcedo *et al*, 2018) proved to be poor predictors of immune responses following AAV gene delivery to the eye. However, a more extensive assessment of local cytokine expression patterns could provide additional insights. Besides, patients with retinal diseases express local inflammation cytokines in the retina that have the potential to leak into the periphery which further complicates the sensitivity of cytokines as biomarkers(Tao *et al*, 2024). Therefore, novel immunomonitoring strategies need to be explored to provide a complete and exhaustive view of immunological status which can further aid in the standardization of the application of immunosuppression strategies in clinics.

Inter-individual variability in immune responses has been observed in human clinical trials as well as in NHPs receiving ocular gene therapy. This is expected since even in human monozygotic twins, the immune responses can differ due to T and B cell repertoire(HAVERKORN *et al*, 1975; Pogorelyy *et al*, 2018). However, with our study we demonstrated that even in a syngeneic murine model where most of the factors were kept the same, high inter-individual variability in immune responses is observed. Even though factors like the surgery and operation of the experiments cannot be exactly the same in all the mice, this kind of artificial variation also exists in human and other animal models(Puranen *et al*, 2023). Similar inter-individual variability of immune response has been reported in vaccine and anti-tumor research within syngeneic murine models(Audebert *et al*, 2021; Rotnemer-Golinkin & Ilan, 2022; Leete *et al*, 2022; Sivan *et al*, 2015). The factors and mechanisms impacting the inter-individual variability are still not well understood. One study applied vaccinia virus to activate the T cell response in C57BL/6 mice to follow CD8^+^ T cell dynamics and inter-individual variability was observed in priming efficiency, effector expansion and memory cell generation(Audebert *et al*, 2021). Additionally, another study reported a varied immune response in C57BL/6 mice administered with concanavalin A (ConA), which can induce immune-mediated hepatitis, and measured kinetics of alanine aminotransferase (ALT) levels (a marker of liver damage), emphasizing immune responses vary significantly in timing and magnitude, even among genetically identical subjects(Rotnemer-Golinkin & Ilan, 2022). Besides, a study tracked CD8^+^ cell in CT26 syngeneic murine tumor model to assess anti-PD-1 therapy response and high variability in anti-tumor responses was observed, indicating contribution of the rate of CD8^+^ T cell activation to the inter-individual variability(Leete *et al*, 2022). Another study showed that different intestinal microbiota could impact the immune response in syngeneic mice. Mice that were raised in two different facilities showed differences in CD8^+^ cell priming and accumulation during anti-tumor therapy(Sivan *et al*, 2015). In our study however, all the mice were raised in the same facility, under the same diet and housing conditions, and are expected to have similar intestinal microbiota. Nonetheless, this can be a factor influencing the variability in human and other animal models and merits consideration. Such studies reinforce the idea that inter-individual immune variability is a common phenomenon in immunological contexts beyond gene therapy and further exploration is needed to elucidate the underlying mechanisms.

Animal models provide crucial inputs for gene therapy development and fundamental immunology research. In our study, despite using a syngeneic murine model to eliminate most internal and external variability factors, an inter-individual variability in immune responses was observed. Alternatively, research on cell and organoid levels, which are used in gene therapy transduction (Ramamurthy *et al*, 2022), may assist in understanding more about the factors impacting the variability by focusing for example on innate immune reactivity. Regardless of the systemic collaboration of immune systems, cell lines and organoid models can still provide the insight of the immune sensitivity to specific antigens where attempts to use cell culture to predict immunogenicity have been initiated(Fischer *et al*, 2011; Ozawa *et al*, 2021).

Our findings underscore the need for individualized patient care strategies, highlighting the value of a personalized medicine approach(Zheng *et al*, 2015; Ong *et al*, 2013). The understanding and surveillance of individual immune response variations could inform personalized decisions about dosing, treatment selection, duration, and immunosuppressive regimens. The contribution of artificial intelligence (AI) in biology has been expanded widely during the last decade, which allows meta-data analysis and predictions. Multiple studies have used AI in cancer immunotherapy and different AI models have been trained to predict not only various immune signatures but also direct immunotherapy responses(Yang *et al*, 2022). Despite the existence of predictive tools of the immune response in specific cells or signaling pathways(Penhaskashi *et al*, 2023; Chandler *et al*, 2022), those have not been adapted to the context of gene therapy, and challenges in predicting immune responses in this field still largely persist. Current limitations in our understanding of immune system complexity may prevent AI from generating accurate predictions.

In humans and complex animal models like NHPs, inter-individual differences in immune responses are expected and attempts are made to minimize their impact on safety and efficacy (immunosuppression strategies) or to resolve symptoms (anti-inflammation strategies), with little to no information on the underlying mechanisms. Our goal with the present study was to identify the most pertinent parameter or a combination of parameters that can be reliably used to predict, follow-up and manage immune responses post-therapy. However, even in syngeneic mice receiving the same treatments we observed high variability in immune responses. Few correlations were observed among local transgene expression, local inflammation, and systemic immune responses revealing the limitation of the current immunomonitoring strategies in ocular gene therapy clinical trials. Our study reinforces two crucial points – the first one being that the current immunomonitoring strategies can hardly be used to infer on the ongoing immune response and adapt the immunosuppressive regimens. Efforts should be focused on the understanding of the underlying mechanisms leading to the individual differences observed in the immune response. Second, in the therapeutic context patients may benefit from ‘personalized therapies’ that are designed taking into consideration their own unique immune profiles.

### Limitations of the study

In this study, a limited range of immune parameters were evaluated concerning local inflammation, systemic cytokines and systemic anti-transgene product and anti-capsid humoral and cellular immune response. These outputs which correspond to those mostly used in clinical immunomonitoring may not reflect the complex individual immune responses after AAV subretinal injection. Therefore, a more thorough analysis of local inflammation or the systemic immune response using alternative techniques can strengthen the conclusions regarding immune response variability while also identifying new potential biomarkers and immunomonitoring strategies. Besides, all the experiments were performed based on a controlled syngeneic murine model with the same age and gender. Our results should be confirmed in other syngeneic murine models to exclude the potential impact of the model, and possibly with additional conditions such as the animals’ age, more AAV serotypes.

## MATERIALS AND METHODS

### Reagents and Tools table

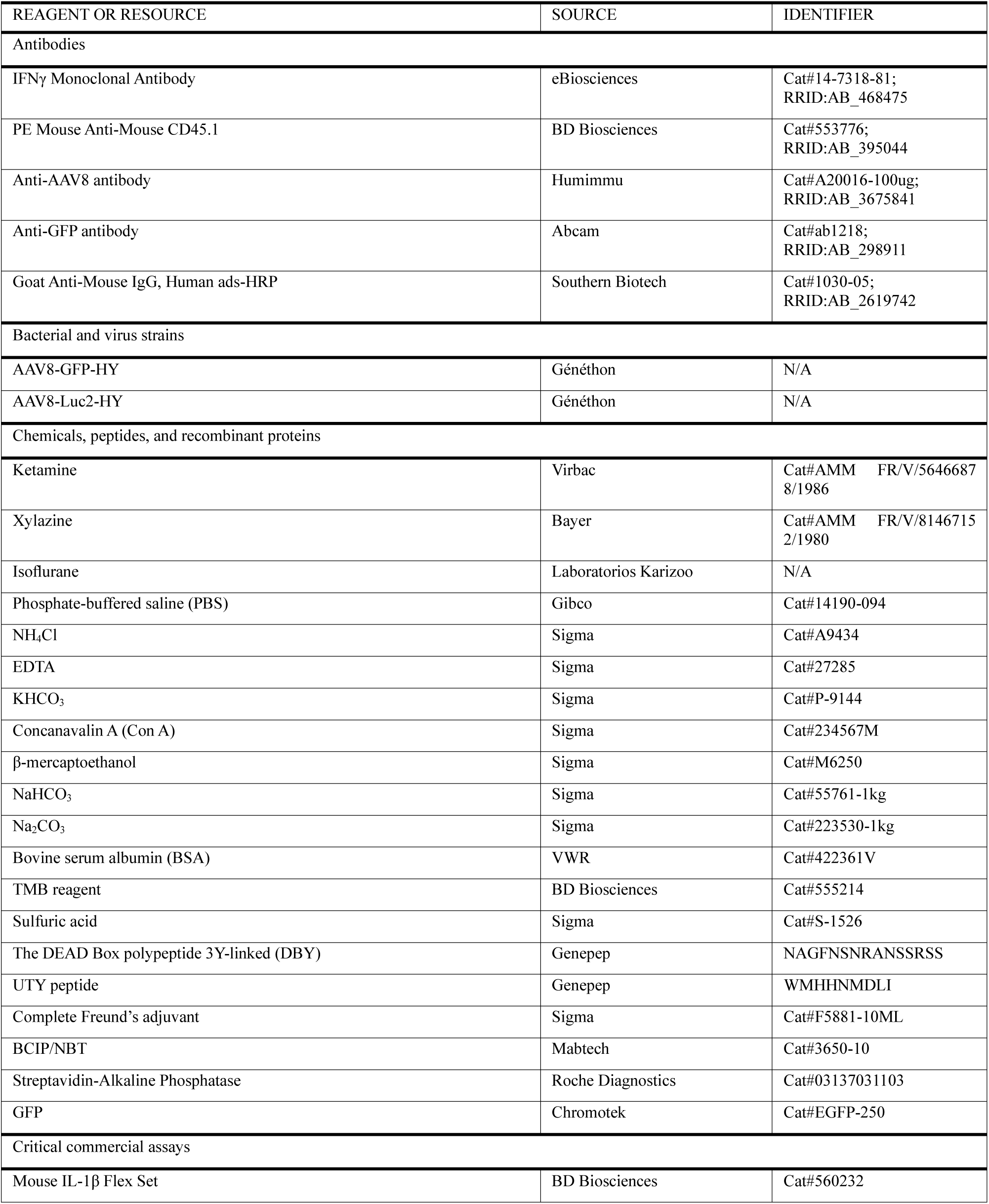
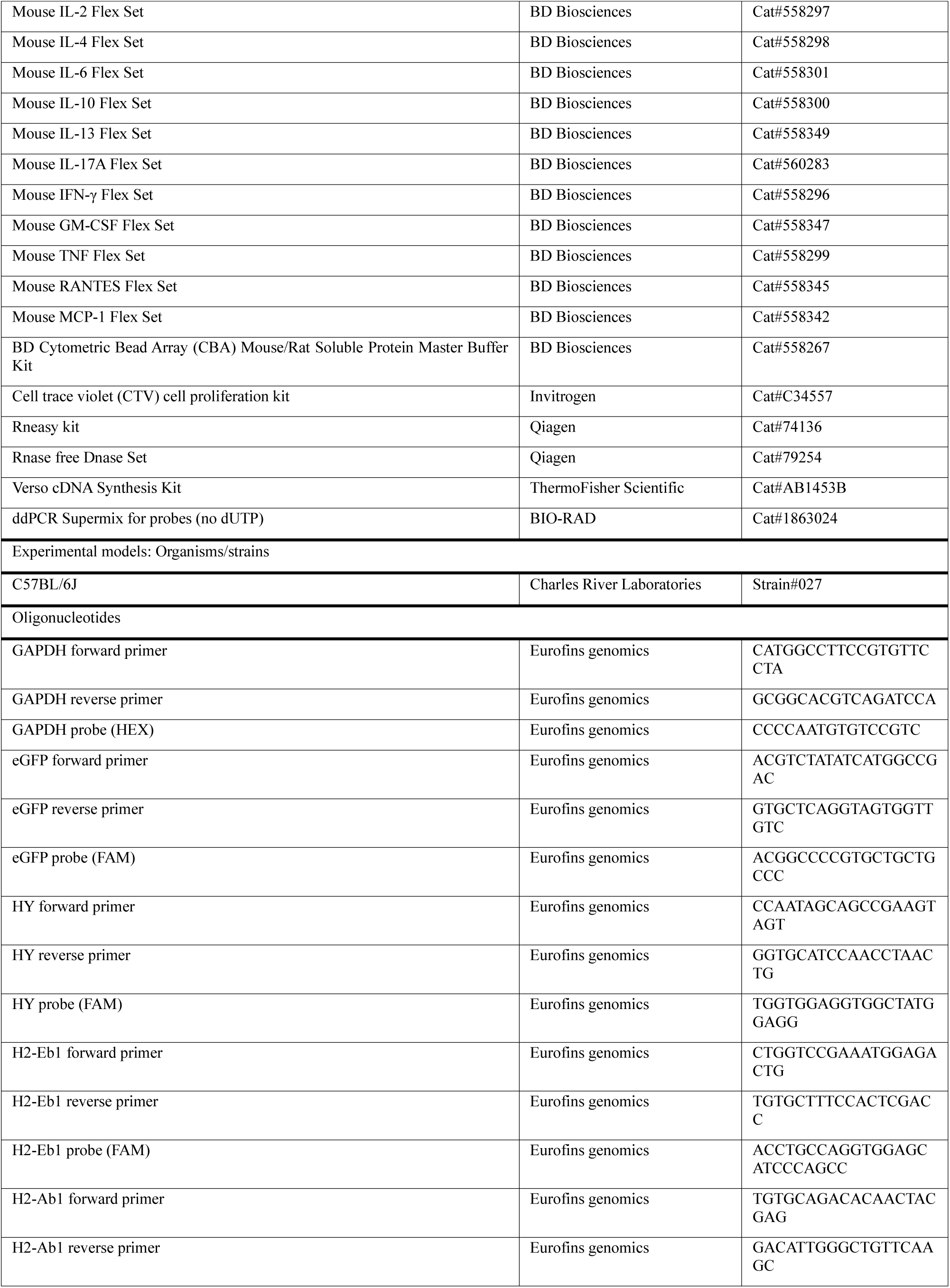
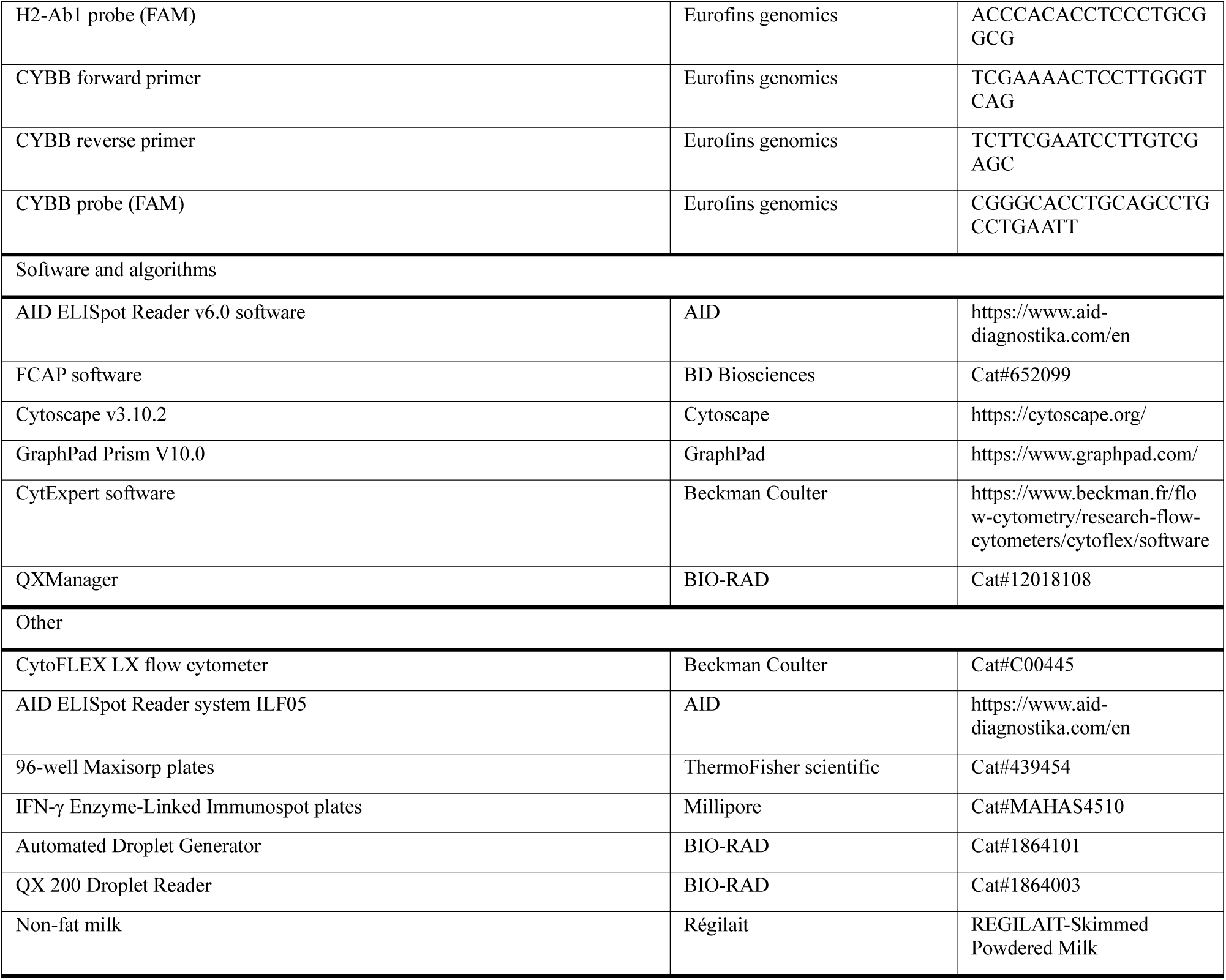

### Methods and protocols

#### Clinical data and NHP data analysis

Immunomonitoring data from the clinical trial (NCT02064569) that includes 15 patients with LHON carrying the G11778A-ND4 mutation was analyzed. These patients were divided into 4 dose cohorts (9×10^9^, 3×10^10^, 9×10^10^, and 1.8×10^11^ vg per eye) and each cohort received an intravitreal injection. Ocular inflammation, anti-AAV2 TAb, anti-AAV2 NAb and anti-AAV2 cellular immune response were quantified, and details were described previously(Bouquet *et al*, 2019). In brief, a composite global OIS was determined according to SUN classification. Anti-AAV2 TAb were determined by ELISA from human serum samples prediluted at 1:50. NAb assay was used to measure anti-AAV2 NAb with HEK293 cells cultured with an rAAV2 expressing luciferase under the control of the cytomegalovirus promoter (8×10^7^vg per well) and either alone or serial fold dilutions of human serum samples. The half maximal inhibitory concentration was determined using the intercept at 50% of the regression curve and expressed as a dilution factor. Peripheral blood mononuclear cells (PBMCs) were isolated and an IFNγ ELISpot assay was performed to measure cellular immune response against AAV2 antigens. Angle sector diagrams for patients were generated based on the FC of anti-AAV2 TAb and NAb, maximal OIS and anti-AAV2 cellular immune response which had been normalized with the highest value corresponding to each aspect.

Immunomonitoring data from 8 NHPs was analyzed. NHPs received an intravitreal injection of 1×10^9^ vg AAV2.7m8 in both eyes. Ocular inflammation, anti-AAV2 TAb and Nab were evaluated and details were described previously(Ail *et al*, 2022). In brief, anti-AAV2 TAb were determined by ELISA from NHP serum samples prediluted at 1:100. NAb assay for AAV2 was performed using HEK293T cells cultured with an rAAV2 expressing luciferase under the control of the cytomegalovirus promoter (at a multiplicity of infection (MOI) of 6,400 per well) and either alone or serial fold dilutions of NHP serum samples. The half maximal inhibitory concentration was determined using the intercept at 50% of the regression curve and expressed as a dilution factor. A Spectralis HRA + OCT system was used to acquire OCT images. SUN classification was applied for grading the “Anterior chamber cells”. For grading the “Vitreous cells”, the NIH grading system is used. The British Medical Journal (BMJ) grading system is used for grading the “Posterior Uveitis”.

#### Murine model

Wild-type 6- to 8-week-old female C57BL/6J mice (H-2^b^) were purchased from Charles River Laboratories (L’Arbresle, France). Animals were anesthetized either by intraperitoneal injection of 120 mg/kg ketamine (Virbac, Carros, France) and 6 mg/kg xylazine (Bayer, Lyon, France) or by inhalation of isoflurane (Baxter, Guyancourt, France). They were euthanized by cervical elongation. All mice were housed, cared for, and handled in accordance with the European Union guidelines and with the approval of the local research ethics committee (CEEA-51 Ethics Committee in Animal Experimentation, Evry, France; authorization number 2015102117539948).

#### AAV vectors

AAV8-GFP-HY and AAV8-Luc2-HY vectors were produced by Généthon in Evry (France) using the tritransfection technique in 293T cells cultured in roller bottles(Liu *et al*, 2003). Transgenes were under the ubiquitous Phosphoglycerate kinase (PGK) promoter. HY is a male antigen which is immunogenic in female mice. AAV vectors were purified by cesium chloride gradients centrifugation, and vector titters were determined by qPCR. Endotoxin levels were below 6 E.U/mL.

#### Peptides

The DEAD Box polypeptide 3Y-linked (DBY) and Ubiquitously Transcribed tetratricopeptide repeat gene Y-linked (UTY) peptides, NAGFNSNRANSSRSS and WMHHNMDLI, respectively, were synthesized by Genepep (Montpellier, France) and shown to be more than 95% pure. UTY and DBY are immunodominant peptides of the HY antigen, restricted to MHC-I and MHC-II, respectively.

#### Injections in mice

Injections were performed as described previously(Vendomèle *et al*, 2024). Briefly, the right eye was protruded under microscopic visualization, and the sclera was perforated with a 27G beveled needle. A blunt 32G needle set on a 10 μL Hamilton syringe was inserted in the hole and the same volume (2 μL) of PBS, or AAV (5×10vg), was injected into the subretinal space via the vitreous. The quality of the injection was verified by checking the detachment of the retina and the absence of reflux outside the eye.

#### Cell extraction from murine spleen

After euthanasia, cells were extracted from spleen as described previously(Vendomèle *et al*, 2018). Briefly, spleens were removed and crushed with a syringe plunger on a 70-µm filter in 2 mL of RPMI medium. Red cells were lysed by adding ACK buffer (8.29 g/L NH_4_Cl, 0.037 g/L EDTA, and 1 g/L KHCO_3_) for 1 min. Lysis was stopped by addition of complete RPMI medium (10% FBS, 1% penicillin/streptomycin, 1% glutamine, and 50 µM β-mercaptoethanol). After centrifugation, cells were counted and the concentration was adjusted in complete RPMI medium.

#### Murine IFNγ ELISpot assay

IFNγ ELISpot assay was performed as described previously(Vendomèle *et al*, 2018). IFN-γ Enzyme-Linked Immunospot plates (MAHAS4510, Millipore, Molsheim, France) were coated with anti-IFNγ antibody (eBiosciences, San Diego, CA, USA) overnight at +4°C. Stimulation media (complete RPMI), AAV (1×10^11^ vg/mL), UTY (2 µg/mL), DBY (2 µg/mL), UTY + DBY (2 µg/mL), or Concanavalin A (Sigma, Lyon, France) (5 µg/mL) were plated and 5×10^5^ spleen cells/well were added. After 24 h of culture at 37°C, plates were washed and the secretion of IFNγ was revealed with a biotinylated anti-IFNγ antibody (eBiosciences), Streptavidin-Alcalin Phosphatase (Roche Diagnostics, Mannheim, Germany), and BCIP/NBT (Mabtech, Les Ulis, France). Spots were counted with an AID ELISpot Reader system ILF05 and the AID ELISpot Reader v6.0 software. Results are expressed in index where IFNγ secretion of the positive control was set to 100, based on the positive control for anti-HY immune response, in order to compile and to compare results from different experiments.

#### Cytokine titration by multiplex cytometric bead array

Cytokine titration by multiplex cytometric bead array was performed as described previously(Vendomèle *et al*, 2018). Stimulation media (complete RPMI), AAV (1×10^11^ vg/mL), UTY (2 µg/mL), DBY (2 µg/mL), UTY + DBY (2 µg/mL), or Concanavalin A (Sigma, Lyon, France) (5 µg/mL) were plated and 1×10^6^ spleen cells/well were added. After 36 h of culture at 37°C, supernatants from triplicates were pooled and frozen at −80°C until the titration. Cytometric bead arrays were performed with BD Biosciences flex kits (IL-2, IL-4, IL-6, IL-10, IL-13, IL-17A, IFN-γ, GM-CSF, TNFα, and MCP-1). Briefly, capturing bead populations with distinct fluorescence intensities and coated with cytokine-specific capture antibodies were mixed together. Next, 25 µL of the bead mix of beads was distributed and 25 µL of each sample (supernatants) was added. After 1 h of incubation at room temperature, cytokine-specific PE-antibodies were mixed and 25 µL of this mix was added. After 1 h of incubation at room temperature, beads were washed with 1 mL of Wash buffer and data were acquired with an LSRII flow cytometer (BD Biosciences). FCAP software (BD Biosciences) was used for the analysis.

Radar diagrams represent the percentage of cytokine secretion in the different groups based on the maximum of cytokine secretion and were performed with Excel software. The 100% radar scale fits the maximum value of cytokine secretion, and the black area values correspond to the means of these cytokine secretions

#### *In vivo* cell cytotoxicity assay

IVC assay was performed as previously described(Vendomèle *et al*, 2024).

Spleen cells from CD45.1^+^ CD45.2^−^ male (expressing HY antigen) and CD45.1^−^ CD45.2^+^ female (not expressing HY antigen) C57BL/6 wild-type mice were harvested as described above and stained with cell trace violet (CTV) cell proliferation kit (Molecular Probes) in PBS at different concentrations: 2 μM for male and 20 μM for female cells, according to the protocol of the kit. CTV staining level allows tracking separately male, and female transferred cells. A mixture of 3×10^6^ male cells (CTV^low^) and 3×10^6^ female cells (CTV^high^) in 200 μL was injected intravenously in the experimented female C57BL/6 mice (CD45.1^−^ CD45.2^+^ and not expressing HY antigen) at day 17 of the protocol. CD45.1^−^ CD45.2^+^ CTV^high^ female cells are used as a control of cell survival, as they are not targeted by anti-HY immune responses. Three days after injection, blood was harvested, red blood cells were lysed by adding ACK buffer, washed in PBS 1X, and leukocytes were stained for flow cytometry with an anti-CD45.1-PE antibody to analyze the male cell survival *in vivo* (Pharmingen, BD Biosciences). Data were acquired on a CytoFLEX LX flow cytometer (Beckman Coulter) and analyzed with the CytExpert software (Beckman Coulter).

#### Enzyme-linked immunosorbent assay (ELISA)

96-well Maxisorp plates (ThermoFisher scientific) were coated with full AAV8 capsid (5×10^8^ vg/well) or GFP (0.5 μg/well, Chromotek) diluted with coating buffer (0.84% NaHCO_3_, 0.356%Na_2_CO_3_, pH9.5) overnight at 4°C. The wells were emptied and washed with blocking buffer (PBS 1X-6% milk) before incubating the plate with blocking buffer at room temperature for 2 hours. Serial dilutions of primary antibodies (Anti-AAV8 antibody, Hum Immu; Anti-GFP antibody, Abcam) or serum were prepared during the incubation. Primary antibodies were diluted with dilution buffer (PBS 1X-1% BSA) to ensure a gradient on the plate (100ng, 50ng, 25ng, 12.5ng, 6.25ng, 3.2ng, 1.56ng, 0.781ng, 0.391ng, 0.195ng, 0.0975ng for AAV8 and 25ng, 12.5ng, 6.25ng, 3.2ng, 1.56ng, 0.781ng, 0.391ng, 0.195ng for GFP) to calculate the standard curve for the experiment. Murine sera were diluted to 1/1000 with dilution buffer for AAV ELISA and 1/2500 for GFP ELISA. After the incubation, primary antibodies dilution and serum dilution were added in the wells and the plate were incubated for 1 hour in the incubator at 37°C after 3 times washing with washing buffer (PBS 1X-0.05% Tween 20). 1/4000 dilution of secondary antibodies (Goat Anti-Mouse IgG, Southern Biotech) was prepared with dilution buffer. Secondary antibodies were added into each well after washing 3 times and the plate was incubated for 1 hour in 37°C incubator. The TMB reagent (BD Biosciences) and stop solution (1M sulfuric acid) were placed at room temperature 30 minutes before the end of the previous incubation. When the last incubation ended, the plate was washed 3 times with washing buffer. TMB reagent was added and a blue color appeared. Stop solution was added to stop the reaction after 10 minutes. The plate was read at 450nm to get the optical density of each sample.

#### RNA extraction from mouse models retinas and reverse transcription

Total RNA was isolated using RNeasy kit, QIAGEN (ref: 74106). The procedure was performed according to the manufacturer’s specification. The purification included a DNase treatment using the RNase free DNase Set (Qiagen). The yield and purity of the RNA was measured with NanoDrop 8000 spectrometer. Verso cDNA Synthesis Kit (ThermoFisher Scientific) was applied for reverse transcription following the protocol provided.

#### Droplet Digital PCR (ddPCR)

For each gene, a mix of primer and probe was prepared with 18µL of primers forward, 18µL of primers reverse, 5µL of probe (Eurofin genomics, initial concentration of 100mM) and 59µL of H_2_0 MilliQ.11µL of ddPCR Supermix for probes (no dUTP) (Biorad), 1µL from the primers/probes mix prepared previously (the ratio of housekeeping gene and target gene was 1:1), 6µL of water, and 2µL of the cDNA sample from mice (diluted with water to obtain quantities of either 5ng or 2.5ng in the wells) were added in one well. The droplets were generated by the Automated Droplet Generator from BIO-RAD. A PCR was proceeded by the C1000 Touch Thermal Cycler from BIO-RAD, with the program as 10min at 95°C, repetition of 30sec at 94°C and 1min at 58.1°C 40 times, then 10min at 98°C to finalize at 12°C infinite hold. The results were obtained by QX 200 Droplet Reader from BIO-RAD with the software QXManager for the analysis. Copy numbers of the genes were obtained. Copy number of gene of interest was normalized with copy number of housekeeping gene and the ratio was multiplied by 100.

#### Quantification and statistical analysis

Statistical analyses were performed with GraphPad Prism V10.0. Mann-Whitney tests, correlation matrix and principal component analysis were performed. p value <0.05: *, <0.01: **, <0.001: ***, <0.0001: ****. Radar diagrams and angle sector diagrams made by Excel software represent the percentage of measured value in the different groups normalized with the maximal value in the corresponding aspect. Network analysis was performed with Cytoscape v3.10.2. The computation of half maximal inhibitory concentration of NAb (IC_50_) has been performed using R.

## ACKNOWLEDGMENTS

D.R. was supported by CSC (China Scholarship Council, No. 202108070132). This work was supported by Fondation de France (Berthe Fouassier), LabEx LIFESENSES (ANR-10-LABX-65) and IHU FOReSIGHT (ANR-18-IAHU-01), DIM C-BRAINS funded by the Conseil Régional d’Ile-de-France. The authors would like to thank Peggy Sanatine for flow cytometry processing, Jérémie Cosette for advice on data analysis. The authors would like to thank Elena Brazhnikova, Céline Nouvel-Jaillard and the MirCEN NHP facility for assistance with NHP experiments. We would like to thank Melissa Desrosiers and Camille Robert for their technical assistance with the production of plasmids and viral vectors for the NHP experiments. The schematic illustrations were created on BioRender.com.

## AUTHOR CONTRIBUTIONS

Conceptualization, S.F., D.A. and D.R.; methodology, D.R., G.C., J.V., E.C. and A.P.; Investigation, S.F., D.A., and D.R.; writing—original draft, D.A. and D.R.; writing—review & editing, C.V., J.P., D.R., D.A., A.G., D.D., H.S. and S.F.; funding acquisition, A.G., D.D. and G.R.; resources, C.V.; supervision, D.A., and S.F.

## DATA AVAILABILITY

This study includes no data deposited in external repositories.

Further information and requests for resources and reagents should be directed to and will be fulfilled by the lead contact, Sylvain Fisson

Any additional information required to reanalyze the data reported in this work paper is available from the lead contact upon request, Sylvain Fisson

---

**Supplementary Figure 1.**
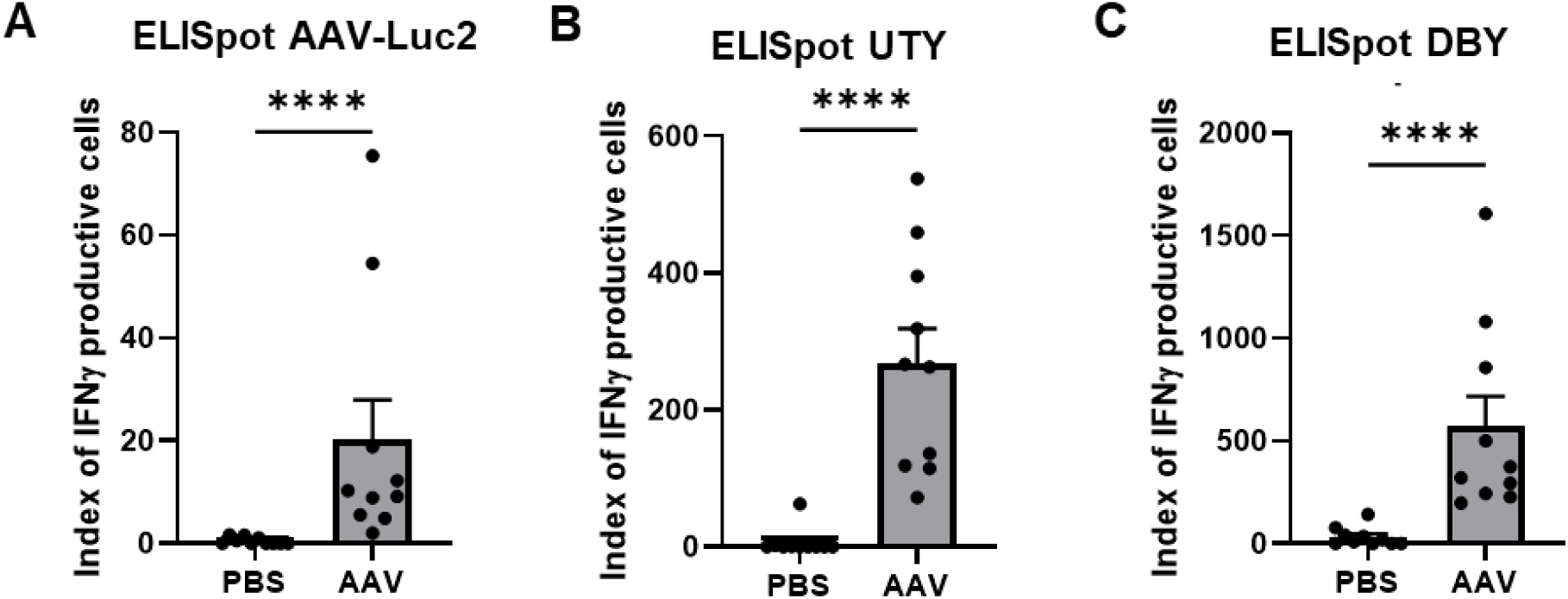
CD4^+^ T cell and CD8^+^ T cell activation by peptides. **A** T cell activation specific to AAV capsid measured by ELISpot assay. **B** CD8^+^ T cell activation specific to UTY peptides. **C** CD4^+^ T cell activation specific to DBY peptides. Data information: Results obtained from 2 independent experiments (n=10 per group). Bars correspond to mean + SEM. *P < 0.05, **P < 0.01, ***P < 0.001, and ****P < 0.0001 with unpaired Mann-Whitney test.

**Supplementary Figure 2.**
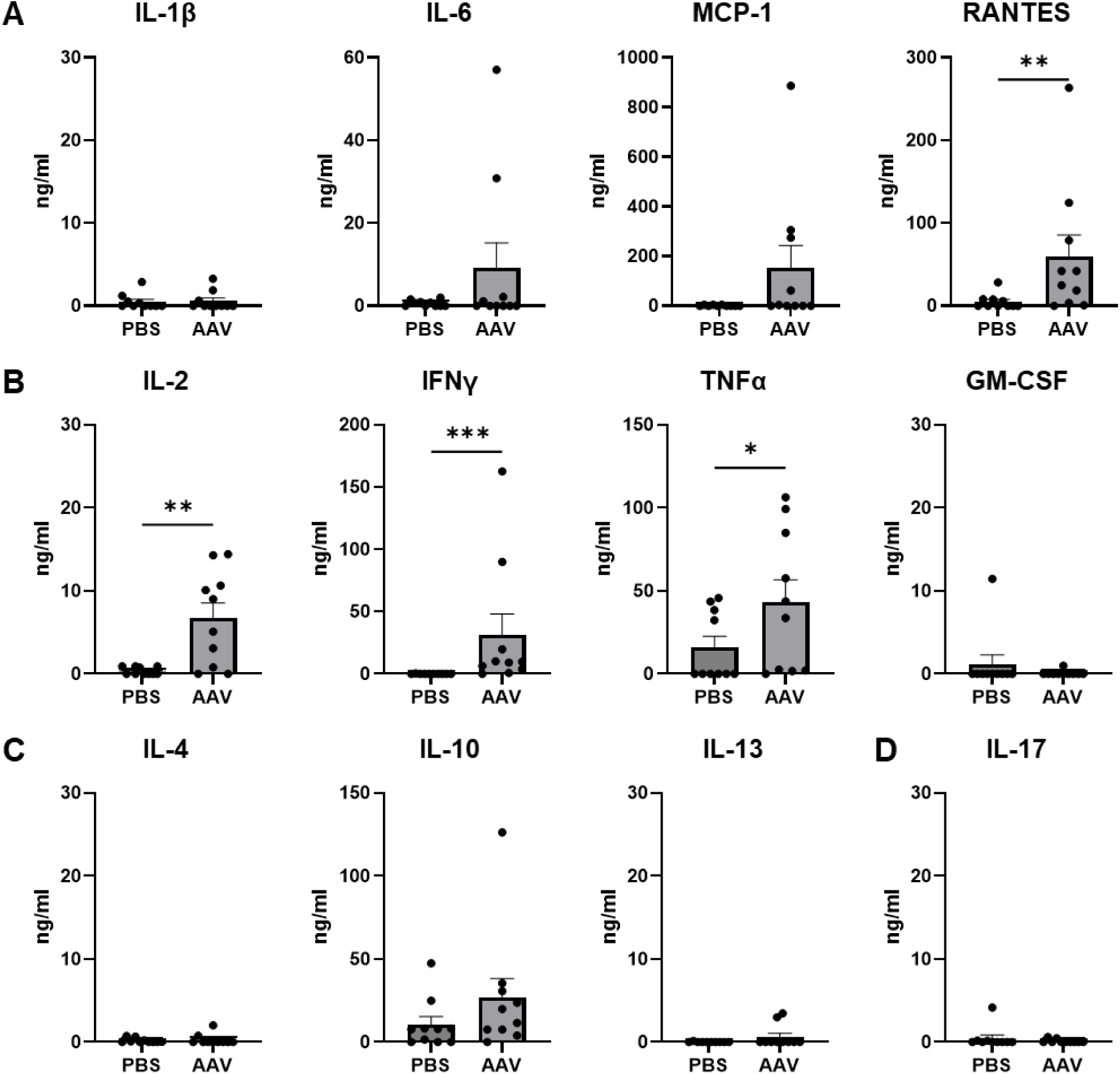
Cytokine secretion in mouse spleen cells 21d post-injection (PI) of AAV-GFP-HY stimulated with AAV *in vitro*. **A-D** Expression of cytokines participating in **(A)** Inflammation and cellular migration (IL-1b, IL-6, MCP-1, RANTES), **(B)** Th1 function (IL-2, IFNγ, TNFα, GM-CSF), **(C)** Th2 function (IL-4, IL-10, IL-13) and **(D)** Th17 function (IL-17). Data information: Results obtained from 2 independent experiments (n=10 per group). Bars correspond to mean + SEM. *P < 0.05, **P < 0.01, ***P < 0.001, and ****P < 0.0001 with unpaired Mann-Whitney test.

**Supplementary Figure 3.**
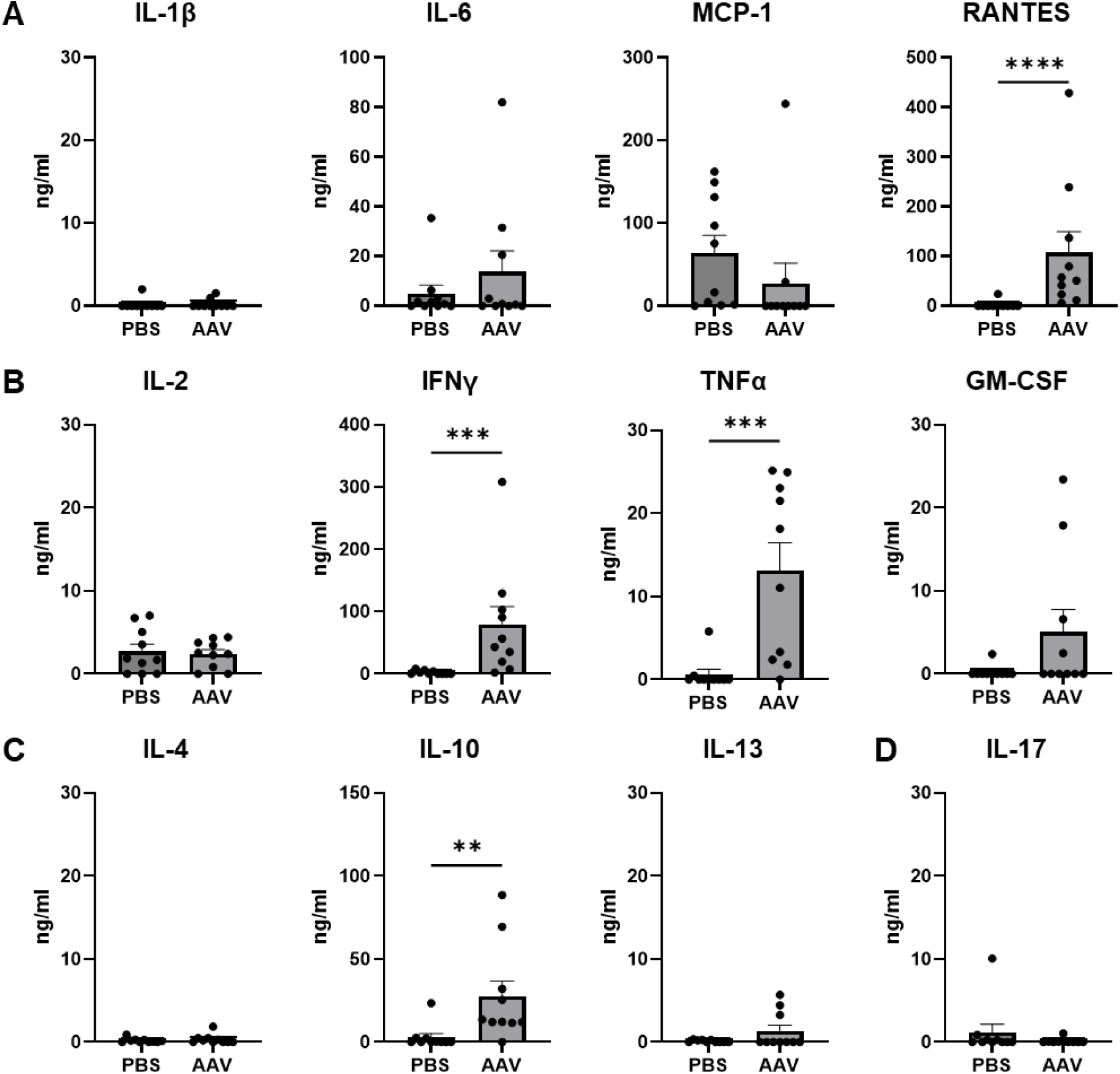
Cytokine secretion in mouse spleen cells 21d post-injection (PI) of AAV-GFP-HY stimulated with HY peptides (pHY) *in vitro*. **A-D** Expression of cytokines participating in **(A)** Inflammation and cellular migration (IL-1b, IL-6, MCP-1, RANTES), **(B)** Th1 function (IL-2, IFNγ, TNFα, GM-CSF), **(C)** Th2 function (IL-4, IL-10, IL-13) and **(D)** Th17 function (IL-17). Data information: Results obtained from 2 independent experiments (n=10 per group). Bars correspond to mean + SEM. *P < 0.05, **P < 0.01, ***P < 0.001, and ****P < 0.0001 with unpaired Mann-Whitney test.

**Supplementary Table 1.** Characteristics of individual immune responses in patients.

| Dose Level,<br>Patient No. | Maximum, OIS | IgG (×10<sup>-3</sup> AU/mL) |  | NAb (IC<sub>50</sub>) |  | Cellular Response |
| --- | --- | --- | --- | --- | --- | --- |
|  |  | Baseline | Maximum Value | Baseline | Maximum Value |  |
| 9E9 vg |  |  |  |  |  |  |
| 001 | 0 | 7115 | 7565 | 1 440 | 3050 | Negative |
| 003 | 0.5 | 3520 | 14895 | 520 | 3650 | Negative |
| 005 | 0 | 1903 | 5355 | 360 | 3400 | Positive |
| 3E10 vg |  |  |  |  |  |  |
| 006 | 1 | 155 | 10665 | 23 | 2200 | Negative |
| 007 | 1.5 | 33 | 336 | 0 | 40 | Negative |
| 008 | 0.5 | 536 | 2585 | 48 | 488 | Negative |
| 9E10 vg |  |  |  |  |  |  |
| 009 | 5 | 2110 | 14240 | 420 | 3200 | Negative |
| 011 | 1.5 | 13 | 1148 | 0 | 120 | Negative |
| 012 | 1 | 32 | 177 | 0 | 60 | Negative |
| 017 | 0.5 | 918 | 1215 | 228 | 396 | Negative |
| 018 | 9.5 | 10295 | 67240 | 2850 | 40 000 | Positive |
| 019 | 1.5 | 26 | 78 | 0 | 24 | Negative |
| 1.8E11 vg |  |  |  |  |  |  |
| 013 | 1.5 | 46 | 760 | 0 | 68 | Negative |
| 014 | 1 | 4952 | 18010 | 630 | 8400 | Negative |
| 015 | 0.5 | 155 | 1229 | 0 | 220 | Negative |

AU, arbitrary unit; IC<sub>50</sub>, half maximal inhibitory concentration; NAb, neutralizing antibody; OI, ocular inflammation; OIS, ocular inflammation score; vg, viral genome.

**Supplementary Table 2.** Characteristics of individual immune responses in NHPs.

| No. | Eye | Anti-AAV2 TAB (%) |  | Anti-AAV2 NAb (IC<sub>50</sub>) |  | Inflammation at Month 1 |  |  |  |
| --- | --- | --- | --- | --- | --- | --- | --- | --- | --- |
|  |  | T0 | T2 | T0 | T2 | ACC | ACF | VH | VC |
| NHP1 | LE | 100 | 453.86 | 23.64 | 443.83 | 3 | 1 | 0 | 1 |
|  | RE |  |  |  |  | 1 | 0 | 0 | 1 |
| NHP2 | LE | 100 | 307.03 | 3.35 | 487.05 | 0.5 | 0 | 0 | 1 |
|  | RE |  |  |  |  | 0.5 | 0 | 0 | 1 |
| NHP3 | LE | 100 | 537.08 | 3.56 | 159.55 | 0 | 0 | 0 | 1 |
|  | RE |  |  |  |  | 0.5 | 0 | 0 | 1 |
| NHP4 | LE | 100 | 408.2 | 1.57 | 309.11 | 1 | 0 | 0 | 1 |
|  | RE |  |  |  |  | 1 | 0 | 0 | 1 |
| NHP5 | LE | 100 | 142.86 | 3.16 | 17.7 | 1 | 0 | 0 | 1 |
|  | RE |  |  |  |  | 1 | 0 | 0 | 1 |
| NHP6 | LE | 100 | 383.36 | 7.94 | 247.3 | 0.5 | 0 | 0 | 1 |
|  | RE |  |  |  |  | 1 | 0 | 0 | 1 |
| NHP7 | LE | 100 | 366.88 | 8.74 | 432.64 | 0.5 | 0 | 0 | 1 |
|  | RE |  |  |  |  | 0.5 | 0 | 0 | 1 |
| NHP8 | LE | 100 | 230 | 100 | 181.75 | 1 | 0 | 0 | 1 |
|  | RE |  |  |  |  | 1 | 0 | 0 | 1 |

TAB, total antibody; NAb, neutralizing antibody; IC<sub>50</sub>, half maximal inhibitory concentration; ACC, anterior chamber cell; ACF, anterior chamber flare; VH, vitreous haze; VC, vitreous cell; LE, left eye; RE, right eye.

**Supplementary Table 3.**
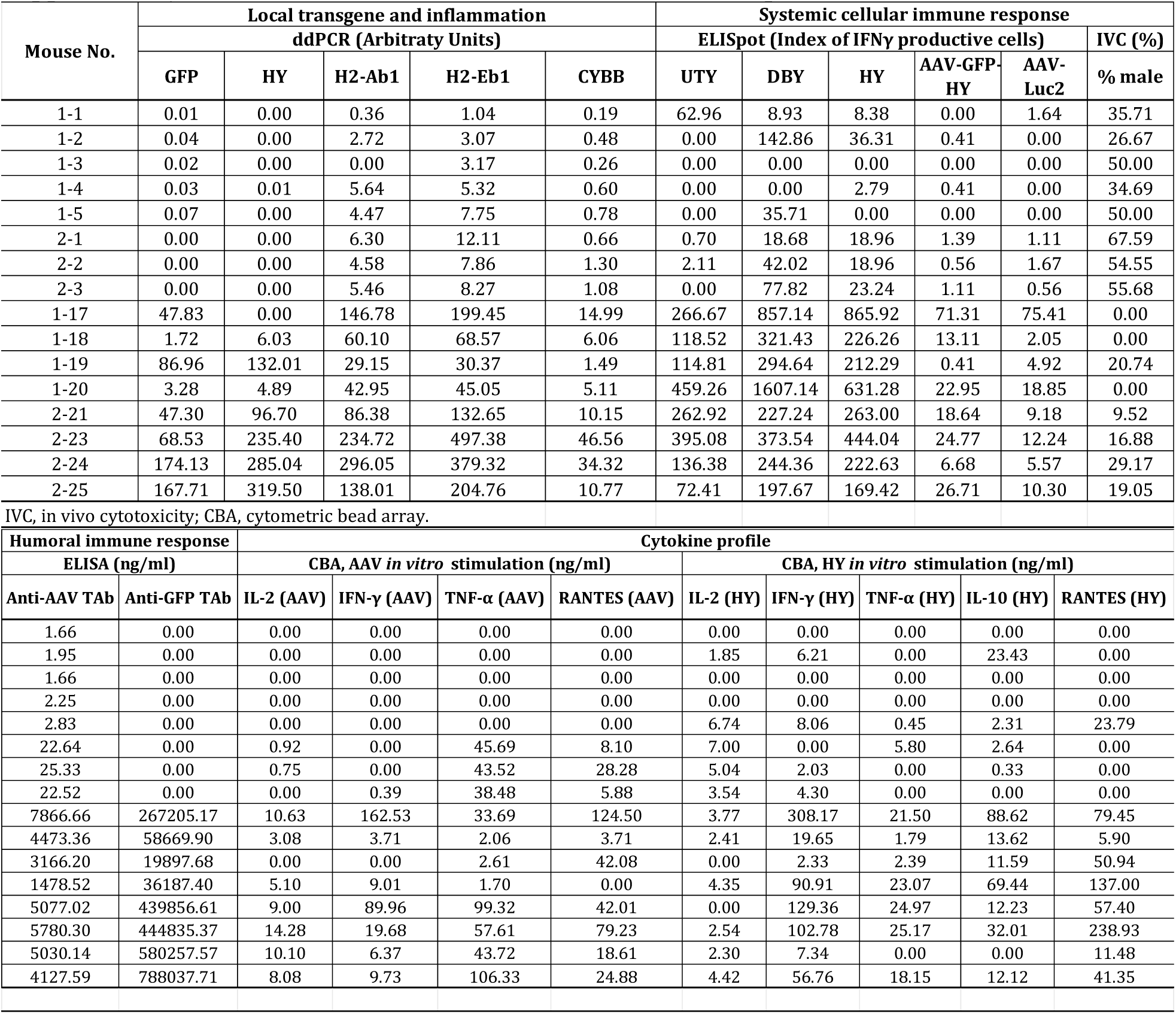
Characteristics of individual immune responses in mice.

| Mouse No. | Local transgene and inflammation |  |  |  |  | Systemic cellular immune response |  |  |  |  |  |
| --- | --- | --- | --- | --- | --- | --- | --- | --- | --- | --- | --- |
| | ddPCR (Arbitraty Units) | | | | | ELISpot (Index of IFNγ productive cells) | | | | | IVC (%) |
|  | GFP | HY | H2-Ab1 | H2-Eb1 | CYBB | UTY | DBY | HY | AAV-GFP-HY | AAV-Luc2 | % male |
| 1-1 | 0.01 | 0.00 | 0.36 | 1.04 | 0.19 | 62.96 | 8.93 | 8.38 | 0.00 | 1.64 | 35.71 |
| 1-2 | 0.04 | 0.00 | 2.72 | 3.07 | 0.48 | 0.00 | 142.86 | 36.31 | 0.41 | 0.00 | 26.67 |
| 1-3 | 0.02 | 0.00 | 0.00 | 3.17 | 0.26 | 0.00 | 0.00 | 0.00 | 0.00 | 0.00 | 50.00 |
| 1-4 | 0.03 | 0.01 | 5.64 | 5.32 | 0.60 | 0.00 | 0.00 | 2.79 | 0.41 | 0.00 | 34.69 |
| 1-5 | 0.07 | 0.00 | 4.47 | 7.75 | 0.78 | 0.00 | 35.71 | 0.00 | 0.00 | 0.00 | 50.00 |
| 2-1 | 0.00 | 0.00 | 6.30 | 12.11 | 0.66 | 0.70 | 18.68 | 18.96 | 1.39 | 1.11 | 67.59 |
| 2-2 | 0.00 | 0.00 | 4.58 | 7.86 | 1.30 | 2.11 | 42.02 | 18.96 | 0.56 | 1.67 | 54.55 |
| 2-3 | 0.00 | 0.00 | 5.46 | 8.27 | 1.08 | 0.00 | 77.82 | 23.24 | 1.11 | 0.56 | 55.68 |
| 1-17 | 47.83 | 0.00 | 146.78 | 199.45 | 14.99 | 266.67 | 857.14 | 865.92 | 71.31 | 75.41 | 0.00 |
| 1-18 | 1.72 | 6.03 | 60.10 | 68.57 | 6.06 | 118.52 | 321.43 | 226.26 | 13.11 | 2.05 | 0.00 |
| 1-19 | 86.96 | 132.01 | 29.15 | 30.37 | 1.49 | 114.81 | 294.64 | 212.29 | 0.41 | 4.92 | 20.74 |
| 1-20 | 3.28 | 4.89 | 42.95 | 45.05 | 5.11 | 459.26 | 1607.14 | 631.28 | 22.95 | 18.85 | 0.00 |
| 2-21 | 47.30 | 96.70 | 86.38 | 132.65 | 10.15 | 262.92 | 227.24 | 263.00 | 18.64 | 9.18 | 9.52 |
| 2-23 | 68.53 | 235.40 | 234.72 | 497.38 | 46.56 | 395.08 | 373.54 | 444.04 | 24.77 | 12.24 | 16.88 |
| 2-24 | 174.13 | 285.04 | 296.05 | 379.32 | 34.32 | 136.38 | 244.36 | 222.63 | 6.68 | 5.57 | 29.17 |
| 2-25 | 167.71 | 319.50 | 138.01 | 204.76 | 10.77 | 72.41 | 197.67 | 169.42 | 26.71 | 10.30 | 19.05 |
| IVC, in vivo cytotoxicity; CBA, cytometric bead array. |  |  |  |  |  |  |  |  |  |  |  |
| Humoral immune response |  | Cytokine profile |  |  |  |  |  |  |  |  |  |
| ELISA (ng/ml) |  | CBA, AAV *in vitro* stimulation (ng/ml) |  |  |  | CBA, HY *in vitro* stimulation (ng/ml) |  |  |  |  |  |
| Anti-AAV Tab | Anti-GFP Tab | IL-2 (AAV) | IFN-γ (AAV) | TNF-α (AAV) | RANTES (AAV) | IL-2 (HY) | IFN-γ (HY) | TNF-α (HY) | IL-10 (HY) | RANTES (HY) | |
| 1.66 | 0.00 | 0.00 | 0.00 | 0.00 | 0.00 | 0.00 | 0.00 | 0.00 | 0.00 | 0.00 |  |
| 1.95 | 0.00 | 0.00 | 0.00 | 0.00 | 0.00 | 1.85 | 6.21 | 0.00 | 23.43 | 0.00 |  |
| 1.66 | 0.00 | 0.00 | 0.00 | 0.00 | 0.00 | 0.00 | 0.00 | 0.00 | 0.00 | 0.00 |  |
| 2.25 | 0.00 | 0.00 | 0.00 | 0.00 | 0.00 | 0.00 | 0.00 | 0.00 | 0.00 | 0.00 |  |
| 2.83 | 0.00 | 0.00 | 0.00 | 0.00 | 0.00 | 6.74 | 8.06 | 0.45 | 2.31 | 23.79 |  |
| 22.64 | 0.00 | 0.92 | 0.00 | 45.69 | 8.10 | 7.00 | 0.00 | 5.80 | 2.64 | 0.00 |  |
| 25.33 | 0.00 | 0.75 | 0.00 | 43.52 | 28.28 | 5.04 | 2.03 | 0.00 | 0.33 | 0.00 |  |
| 22.52 | 0.00 | 0.00 | 0.39 | 38.48 | 5.88 | 3.54 | 4.30 | 0.00 | 0.00 | 0.00 |  |
| 7866.66 | 267205.17 | 10.63 | 162.53 | 33.69 | 124.50 | 3.77 | 308.17 | 21.50 | 88.62 | 79.45 |  |
| 4473.36 | 58669.90 | 3.08 | 3.71 | 2.06 | 3.71 | 2.41 | 19.65 | 1.79 | 13.62 | 5.90 |  |
| 3166.20 | 19897.68 | 0.00 | 0.00 | 2.61 | 42.08 | 0.00 | 2.33 | 2.39 | 11.59 | 50.94 |  |
| 1478.52 | 36187.40 | 5.10 | 9.01 | 1.70 | 0.00 | 4.35 | 90.91 | 23.07 | 69.44 | 137.00 |  |
| 5077.02 | 439856.61 | 9.00 | 89.96 | 99.32 | 42.01 | 0.00 | 129.36 | 24.97 | 12.23 | 57.40 |  |
| 5780.30 | 444835.37 | 14.28 | 19.68 | 57.61 | 79.23 | 2.54 | 102.78 | 25.17 | 32.01 | 238.93 |  |
| 5030.14 | 580257.57 | 10.10 | 6.37 | 43.72 | 18.61 | 2.30 | 7.34 | 0.00 | 0.00 | 11.48 |  |
| 4127.59 | 788037.71 | 8.08 | 9.73 | 106.33 | 24.88 | 4.42 | 56.76 | 18.15 | 12.12 | 41.35 |  |

